# Realistic retinal modeling unravels the differential role of excitation and inhibition to starburst amacrine cells in direction selectivity

**DOI:** 10.1101/2021.06.22.449374

**Authors:** Elishai Ezra-Tsur, Oren Amsalem, Lea Ankri, Pritish Patil, Idan Segev, Michal Rivlin-Etzion

## Abstract

Retinal direction-selectivity originates in starburst amacrine cells (SACs), which display a centrifugal preference, responding with greater depolarization to a stimulus expanding from soma to dendrites than to a collapsing stimulus. Various mechanisms were hypothesized to underlie SAC centrifugal preference, but dissociating them is experimentally challenging and the mechanisms remain debatable. To address this issue, we developed the Retinal Stimulation Modeling Environment (RSME), a multifaceted data-driven retinal model that encompasses detailed neuronal morphology and biophysical properties, retina-tailored connectivity scheme and visual input. Using a genetic algorithm, we demonstrated that spatiotemporally diverse excitatory inputs – sustained in the proximal and transient in the distal processes – are sufficient to generate experimentally validated centrifugal preference in a single SAC. Reversing these input kinetics did not produce any centrifugal-preferring SAC. We then explored the contribution of SAC-SAC inhibitory connections in establishing the centrifugal preference. SAC inhibitory network enhanced the centrifugal preference, but failed to generate it in its absence. Embedding a direction selective ganglion cell (DSGC) in a SAC network showed that the known SAC-DSGC asymmetric connectivity by itself produces direction selectivity. Still, this selectivity is sharpened in a centrifugal-preferring SAC network. Finally, we use RSME to demonstrate the contribution of SAC-SAC inhibitory connections in mediating direction selectivity and recapitulate recent experimental findings. Thus, using RSME, we obtained a comprehensive mechanistic understanding of SACs’ centrifugal preference and its contribution to direction selectivity.

## Introduction

Retinal direction selectivity emerges in direction selective retinal ganglion cells (DSGCs), which strongly respond to motion in one (preferred) direction and weakly to motion in the opposite (null) direction (**Figure 1a**) (Wei 2018; Vaney et al. 2012; Mauss et al. 2017; Borst and Euler 2011). The key mechanism for generating direction selectivity in DSGCs is asymmetric GABAergic inhibition from starburst amacrine cells (SACs) (Wei et al. 2011; Yonehara et al. 2011; Fried et al. 2002). This asymmetry is achieved by asymmetric wiring from SACs to DSGCs (Briggman et al. 2011) combined with the centrifugal (CF) preference of SAC processes (SAC dendrites and axons are synonymous and called processes): SAC processes respond more strongly to motion away from the cell soma (centrifugal) than towards cell soma (centripetal) (**Figure 1b, c**; Euler et al. 2002; Vlasits et al. 2016; Ding et al. 2016; Ankri et al. 2020).

**Figure 1.**
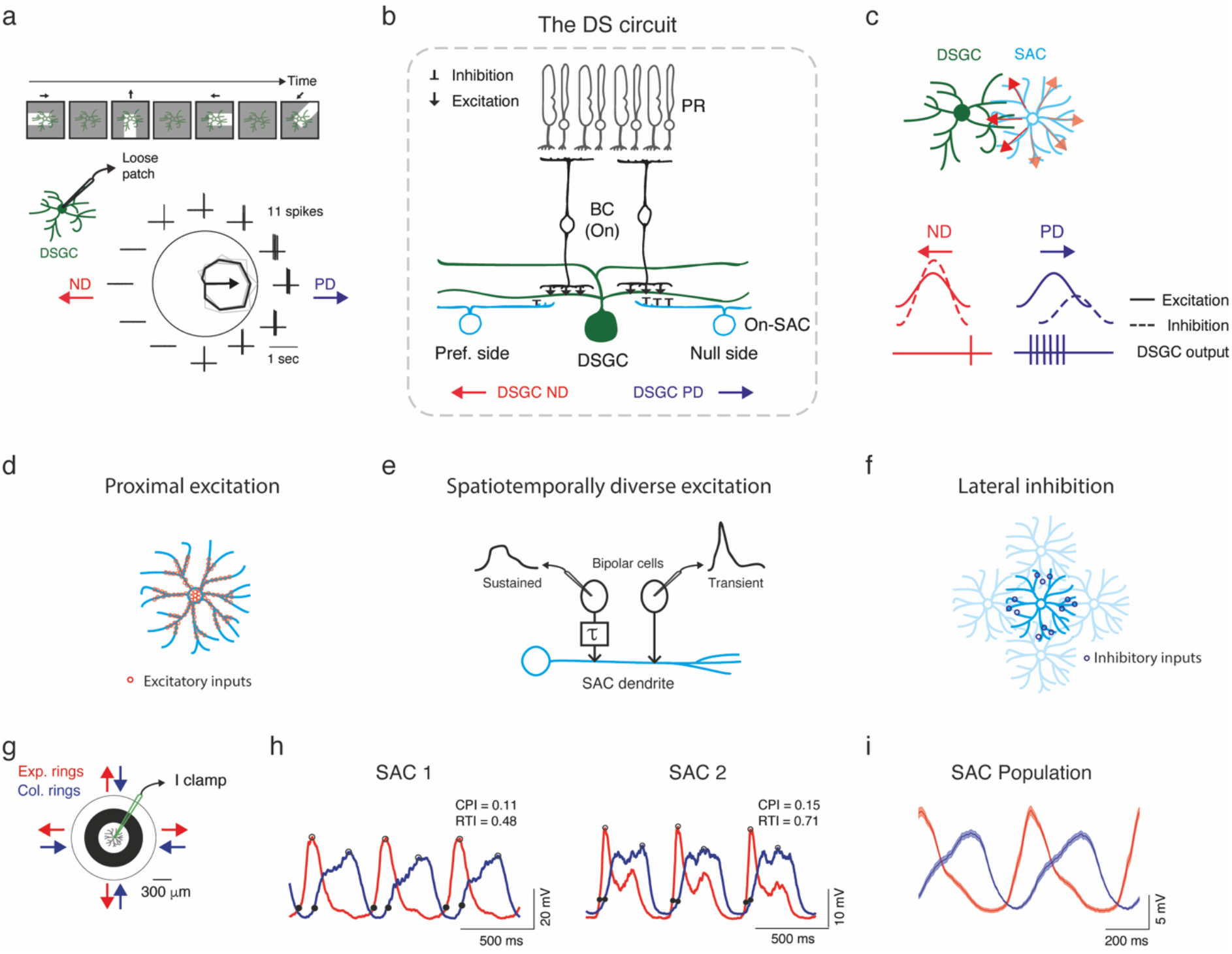
Mechanisms for direction selectivity and centrifugal preference in electrophysiologically recorded SAC. **a.** Example directional tuning of a DSGC. Top: An illustration of a moving bar stimulation. Bottom: Spiking activity recorded in loose-patch mode. The polar plot represents the mean spike count (black) in response to the leading edge of a white bar (On response) moving in 12 directions; grey lines represent single repetitions (4 in total); the arrow represents the preferred direction. The surrounding traces depict one representative recording trace. **b.** A cross-section of the direction selective circuit. Only inputs to DSGCs from the On layer are illustrated for simplicity. DSGCs receive excitatory inputs (↓) from bipolar cells and inhibitory inputs (⊥) from SACs. SACs on the null side form stronger inhibitory connections than SACs on the preferred side. **c.** Schematic of a DSGC innervated by a null-side SAC, top view. SAC processes respond with greater depolarization to centrifugal motion (red arrows), corresponding to the DSGC’s null direction. Bottom: illustration of the excitatory and inhibitory inputs to DSGC during preferred and null motion, and their resulting spiking activity. **d-f.** Illustration of the mechanisms that are thought to underlie SAC CF preference: proximal distribution of excitatory inputs (**d**), differential input kinetics (**e**), and SAC-SAC inhibitory connections (**f**). **g.** Experimental design: current-clamp recordings were performed from On-SACs in response to expanding and collapsing rings centered on the cell soma. **h.** Two examples of SACs voltage response to expanding (red) and collapsing (blue) rings stimulation. Responses to three cycles of the rings are shown. Traces are an average of 5 repetitions. Black filled dots denote the initial response, and empty black dots indicate the peak. **i.** Mean±SEM waveforms of all experimentally recorded SAC responses to expanding and collapsing rings averaged over 1 second (n=27 cells). DSGC: direction selective retinal ganglion cell; SAC: starburst amacrine cell; BC: bipolar cell; PR: photoreceptor; PD: preferred direction; ND: null direction

Different hypotheses have been raised to account for SACs’ CF preference, including their intrinsic properties, inhibitory network connections and their excitatory input distribution (Borst and Euler 2011; Chen and Wei 2018; Demb 2007; Vaney et al. 2012; Wei and Feller 2011; Mauss et al. 2017). The input distribution hypothesis is supported by two recent studies showing that SAC excitatory inputs are confined to its proximal 2/3 of dendritic arbors and skewed away from the distal release sites, an organization that is thought to underlie SAC CF preference (**Figure 1d**; Ding et al. 2016; Vlasits et al. 2016). In addition, physiological and anatomical data have detected different kinetics of the excitatory inputs from bipolar cells to SAC, with more sustained excitation in proximal processes and more transient excitation towards distal processes (Fransen and Borghuis 2017; Kim et al. 2014). The precise spatiotemporal distribution may contribute to SAC CF preference, as only during centrifugal motion the sequential activation of the sustained and transient inputs is effectively integrated (**Figure 1e**). Yet, this spatiotemporal dependence of excitation has not always been observed in the data and its contribution to SAC CF preference remains controversial (Stincic et al. 2016; Ding et al. 2016).

The reciprocal connections between SACs form a dense inhibitory network, which has also been suggested to contribute to SAC CF preference (**Figure 1f**; Morrie and Feller 2018; Lee and Zhou 2006; Fried et al. 2005). However, the role of inhibitory connections between SACs is also unsettled as blocking GABA receptors fails to fully eliminate SACs’ CF responses (Hausselt et al. 2007; Oesch and Taylor 2010; Chen et al. 2016; Hanson et al. 2019). Another recent study suggested that SAC-SAC inhibitory connections mildly affect SAC activity and are more dominant for the computation of direction selectivity in DSGCs under certain stimulus conditions, particularly when the moving stimulus is presented on a noisy background (Chen et al. 2020).

Surprisingly, despite the ample studies dedicated to identifying the source of SAC CF preference, its role in mediating direction selectivity in DSGC is not yet solved. Whereas selective reduction of SAC CF preference was found to decrease direction selectivity (Chen et al. 2016), another study rendered inhibitory inputs to DSGC symmetric (indicating loss of SAC CF preference) and direction selectivity was still maintained (Hanson et al. 2019).

To find the mechanistic balances for SAC CF preference, we developed the Retinal Stimulation Modeling Environment (RSME) framework: a modeling environment for highly detailed, biologically plausible simulations tailored to the exploration of visual processing in retinal circuits. RSME is a versatile framework that supports single cell as well as network modeling. It incorporates detailed morphological and biophysical constraints of each neuron alongside providing efficient modules for retinal mosaic organization, connectivity schemes, synaptic dynamics and graded synaptic release. RSME has a module for generating structured visual stimuli, allowing to assess the responses of the simulated neurons to various light patterns. Using RSME, we aimed to reveal the network mechanisms that can generate a CF preference in simulated SACs that resembles our electrophysiological recordings. We further pushed the neuronal circuits to extreme conditions that are experimentally unfeasible to unfold the role of excitatory input kinetics arrangement and the contribution of SAC-SAC inhibitory connections to SAC CF preference and direction selectivity in DSGC.

## Results

### Electrophysiological recordings reveal SAC CF preference in response to moving rings

We previously demonstrated SAC CF preference using patch-clamp recordings from On-SACs in the isolated mouse retina (Ankri et al. 2020). The electrophysiological SAC recordings presented here combine published and new data. The retina was presented with expanding and collapsing rings centered on the SAC soma, and SAC voltage was recorded in current-clamp mode (**Figure 1g**). In accordance with previous findings (Euler et al. 2002; Vlasits et al. 2016; Ding et al. 2016; Ankri et al. 2020), expanding rings, which generate centrifugal motion, evoked a stronger and faster depolarization response in SACs than collapsing rings, which generate centripetal motion (**Figure 1h**). We used the Centrifugal Preference Index (CPI, see Methods) to assess SAC CF preference based on its response amplitude. CPI values tended to be positive (0.18±0.17, mean ± STD), indicating larger response amplitudes to centrifugal motion. SAC CF preference was also reflected in their response kinetics, with faster responses to centrifugal than to centripetal motion. To quantify the response rise time, we measured the time from the initial response (when the voltage reached 20% of the peak) to peak response (Ankri et al. 2020). We then determined the Rise Time Index (RTI, see Methods) and found it positive (0.33±0.22, mean ± STD). The positive values indicate shorter rise times during centrifugal motion. The average voltage response to expanding and collapsing rings of all experimentally recorded SACs (n=27 cells) further reflects the CF preference in both amplitude and response kinetics (**Figure 1i**; **Figure 4e, g**).

### Modeling Environment

RSME encapsulates NEURON to provide a retina-focused modeling framework in which simulated neurons can be stimulated with visual patterns. RSME comprises a NeuroML-inspired XML-based specification interface (Gleeson et al. 2010) and a dedicated parsing engine, which supports detailed biophysical, morphological, network architecture, and stimulation parameters. It features a set of mechanisms for retinal circuitry related specifications, including (1) graded synaptic transmission-based communication, which is essential as most retinal neurons do not produce action potentials and use graded neurotransmitter release instead; (2) orientation-based rules for synaptic connectivity to accommodate previously reported retinal connectivity patterns (Briggman et al. 2011); (3) arrangement of cells of the same type in a grid, supporting retinal mosaic organization (Heukamp et al. 2020; Masland 2012). Data specification (e.g., number of neurons, synaptic distribution and location) can be visualized with a dedicated module and is processed to generate a NEURON model. RSME also supports iterative execution with varying initial conditions, which can be used for model optimization, as demonstrated below using a genetic algorithm. Results are logged, saved and visualized, and are available for further analysis. A schematic of the software architecture is given in **Figure 2**. Detailed information on RSME architecture can be found in Methods (see **Supplementary Figures 1-3**) and in the project’s GitHub. RSME is an open-source framework available at: https://github.com/NBELab/RSME.

**Figure 2.**
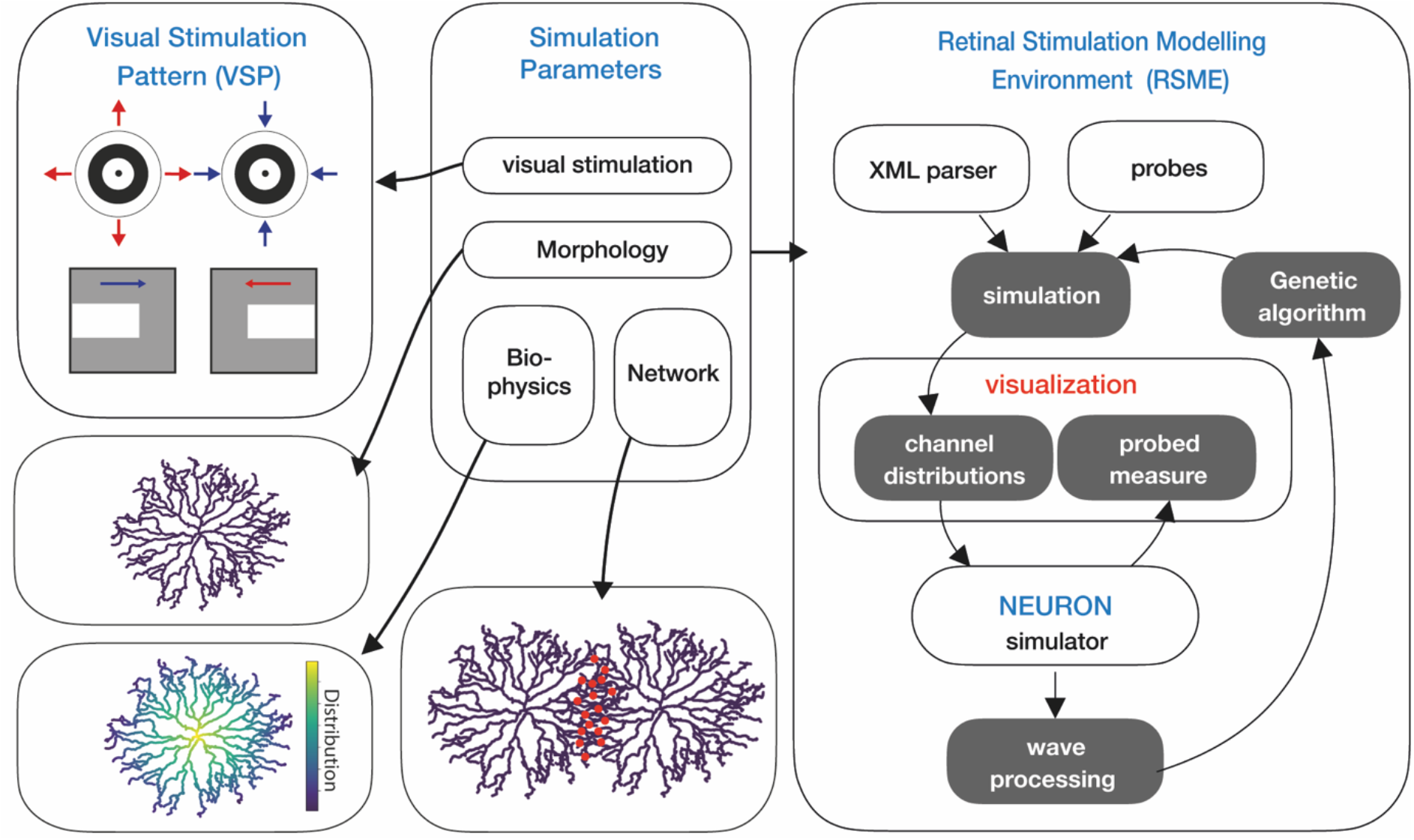
RSME schematic. RSME is comprised of model specifications and a simulation environment. Parameters include the visual stimulation pattern, cellular morphology and biophysics, and network organization. Simulation parameters are parsed and visualized within the modeling environment and used to simulate over NEURON. RSME permits the utilization of signal processing for local optimization (e.g., via a genetic algorithm).

### Simulating a single SAC: SAC’s spatiotemporal excitatory inputs can generate CF preference

We first utilized RSME and a genetic algorithm to test whether a set of synaptic properties can give rise to the CF preference in a single SAC (Ankri et al. 2020; Euler et al. 2002; Fransen and Borghuis 2017). We used the morphology of a reconstructed On-SAC (Ankri et al. 2020) (Neuromorpho.org; ID: NMO_139062), transformed it into a 3D multicompartmental model and specified the passive properties of the cell (**Supplementary Figure 4**). Next, we used a genetic algorithm to scan for a combination of spatial organization and temporal dynamics of excitatory inputs that induce SAC CF preference (**Supplementary Figure 5a-f**). For each set of synaptic inputs, the simulated SAC was presented with expanding and collapsing rings. Centrifugal preference was assessed based on response amplitude (measured as CPI) and rise time (RTI), similar to the experimental data.

Our genetic algorithm explored an 8-dimensional parameter space that controls the distribution and response kinetics of the synapses. Three parameters control the distribution of bipolar cell synapses along the SAC processes according to a sigmoid function, going from denser in the proximal processes to sparser in the distal processes as previously shown (Ding et al. 2016; Vlasits et al. 2016) (**Figure 3a**). The distribution function was parameterized by a proximal density value (synapse probability per 1 μm), a density transition point (the distance from the soma in which input density changed; this point divides the densely innervated proximal processes from the sparsely innervated distal processes), and a distal density value (**Figure 3a**; see *Methods*). Four parameters control the kinetics of bipolar cell synapses, designed using a stochastic vesicle release mechanism (Singer and Diamond 2006). These synapses were regulated by vesicle refiling rate, release probability, a kinetic transition start-point that dictates the location where the synaptic inputs start to change from sustained to transient, and a kinetic transition endpoint that dictates the area where the kinetics remain constant (See *Methods*) The release kinetics were set to shift gradually from more sustained in proximal to more transient in distal synapses (**Figure 3b)** (Fransen and Borghuis 2017; Kim et al. 2014). The eighth parameter controls the conductance of the bipolar cell synapses.

**Figure 3.**
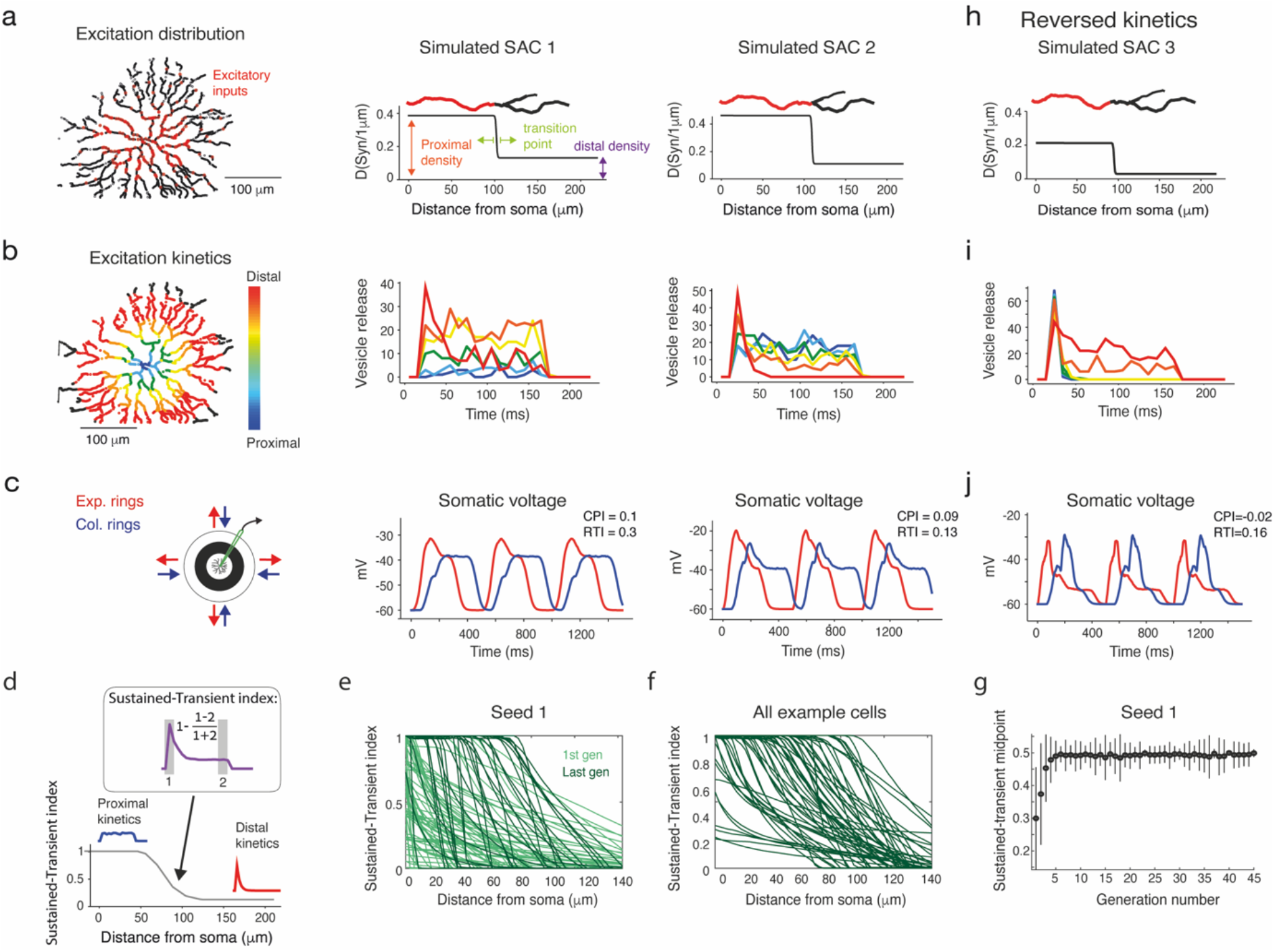
Kinetic properties of excitatory inputs can generate CF preference in simulated SACs. **a.** Left: Reconstruction of a SAC showing the spatially restricted simulated excitatory inputs (red dots). Right: Density of excitatory synapses as a function of distance from soma for two example simulated SACs. The distribution is set by three parameters: proximal synapse density, distal synapse density and the anatomical transition point. For illustration, the transition of color in the example process on top depicts the location of the anatomical transition point. **b.** Left: Illustration of the spatiotemporally diverse excitation distribution, color-coded according to distance from the SAC soma. Right: The kinetics of excitatory inputs in different locations along SAC processes are color-coded by their distance from the cell soma for the two example simulated SACs. **c.** Left: Illustration of the stimulus. Right: The somatic voltage of the two simulated SACs in response to expanding (red) and collapsing (blue) rings. **d**. The sustained-transient index of the excitatory inputs was calculated at each dendritic location based on the input kinetic waveform (see inset and Methods). Values of 1 and 0 indicate completely sustained and transient input kinetics, respectively. **e**. The sustained-transient index as a function of distance from cell soma for all cells in the first generation (light green) and last generation (#45; dark green) of an example simulation seed. **f.** Same as **e**, for all example cells shown in **Figure 4d&f**. **g**. The midpoint of the sustained-transient index for all cells in the example seed as a function of generation. By the 6^th^ generation, the indices tend to span the entire range – from 1 in the proximal to 0 in the distal processes – and the index converges on a value of 0.5. **h-j**. As in **a-c** but for an example SAC with reversed kinetics of the excitatory inputs, changing from transient inputs in the proximal to sustained inputs in the distal processes.

Many different parameter configurations were found to produce a CF preference in SAC, implying that various combinations of anatomical constrains on the excitatory inputs and their kinetics can lead to SAC CF preference. We used six different seeds for the genetic algorithm and found that the results were qualitatively the same, indicating that the convergence is independent of the initial parameters. The set of cells that displayed a clear CF preference, as indicated by Amp_CF_-Amp_CP_≥4, RT_CF_-RT_CP_≥0, and Voltage_score_≥0 (see Methods), is localized to only a subset of the initial distribution, indicating that only a smaller space of parameters produces realistic responses (**Supplementary Figure 6**). Here, the anatomical transition point provided an insight into the anatomical distribution of excitatory inputs that can generate SAC CF preference: as mentioned above, the distribution of inputs from bipolar cell synapses is confined to SAC’s proximal 2/3 dendritic arbors (Ding et al. 2016; Vlasits et al. 2016; Ankri et al. 2020). The genetic algorithm demonstrated that CF preference in SACs could also arise when the transition point is significantly closer to the SAC soma than observed biologically (**Supplementary Figure 6**). Yet, the retina does not implement this solution, probably to maximize the excitatory receptive field area of SACs.

Two example sets of SAC parameters are shown together with the voltage trace produced by RSME (**Figure 3a-c**). In these examples, the distribution of inputs from bipolar cell synapses matched the known anatomical constraints of SAC excitatory inputs. These representative examples demonstrate the similarity of the results of the genetic algorithm and our experimental recordings (compare **Figure 3c** to **Figure 1h**).

We next investigated the contribution of non-homogenous input kinetics in generating SAC CF preference. For this, we first assessed the input kinetics along the SAC process based on the sustained-transient index of the synaptic input as a function of its location (**Figure 3d**). According to the constrains we set, the sustained-transient index decreased with distance from cell soma. However, the dynamics of this decrease differ between the first and the last generations of the genetic algorithm. In the first generation, before the simulation converged to CF-preferring SACs, the decrease in the sustained-transient index was moderate and started already at the very proximal processes. As the algorithm converged and the generations primarily comprised CF-preferring SACs, the decline of the sustained-transient index was steeper and located further from cell soma (**Figure 3e-g**).

Second, we manipulated the input kinetics to follow the reversed logic by running the genetic algorithm once more. Here, transient inputs were confined to the proximal dendrites and became more sustained towards the distal dendrites. All other parameters space remained unchanged. Interestingly, under these reversed-kinetics conditions, the algorithm did not converge and none of the SACs displayed CF preference in this run (**Supplementary Figure 5g-i**). Instead, some SACs revealed a slight centripetal preference (**Figure 3h-j**). Taken together, our simulation results suggest that the distribution of excitatory input kinetics along SAC processes plays a vital role in determining its CF preference.

### Simulating SAC network: SAC-SAC inhibitory connections enhance SAC CF preference

To investigate the contribution of inhibitory connections between SACs to SAC CF preference, we used RSME to create networks of SACs and stimulated them with expanding and collapsing rings, which were centered on the central cell. To form these networks, we manually chose simulated SACs from the results obtained by the genetic algorithm. We restricted our choice to SACs whose set of parameters produced a CF preference (CPI>0) and obeyed the anatomical constraint of a dense excitatory input limited to the inner 1/2-2/3 of their processes (Ding et al. 2016; Vlasits et al. 2016; Ankri et al. 2020) (n=76 SACs). We then replicated each simulated SAC to create a two-layered design, consisting of two overlaid 3×3 and 2×2 grids of SACs (a total of 13 cells, **Figure 4a**). Within this SAC network, inhibitory GABAergic synapses were located at intersecting sections (see Methods). Synapse direction (source/target) was randomly determined. **Figure 4b** illustrates the CF preference of the central SAC as a function of inhibition weight between neighboring SACs in an example SAC network. Inhibition weight was determined by the conductance of the inhibitory synapses. In the absence of inhibition, the cells are independent, and the simulation returns to the initial response of a single SAC given the set of parameters found by the genetic algorithm. As depicted in the simulation example in **Figure 4b**, we found that while SAC inhibitory network only slightly enhanced SAC CF preference in terms of response amplitude, it significantly enhanced CF preference in terms of response kinetics (assessed by CPI and RTI, respectively). To further examine the differential contribution of lateral inhibition and distribution of excitation kinetics to SAC CF preference, we simultaneously shifted two parameters in the network configuration: the inhibition strength and the kinetic transition start-point (where the bipolar cell inputs shift from sustained to transient dynamics). We assessed the CF preference of the example SAC based on its response CPI and RTI (**Figure 4c**). In the absence of inhibitory connections, the CPI and RTI values were low regardless of the transition point and tended to increase with inhibition weight. This increase was more evident for the RTI values, as the CPI values tended to depend both on the inhibition weight and the kinetic transition point. These data imply an interplay of inhibitory and excitatory mechanisms coactivated in the SAC circuit to enhance its CF preference.

**Figure 4.**
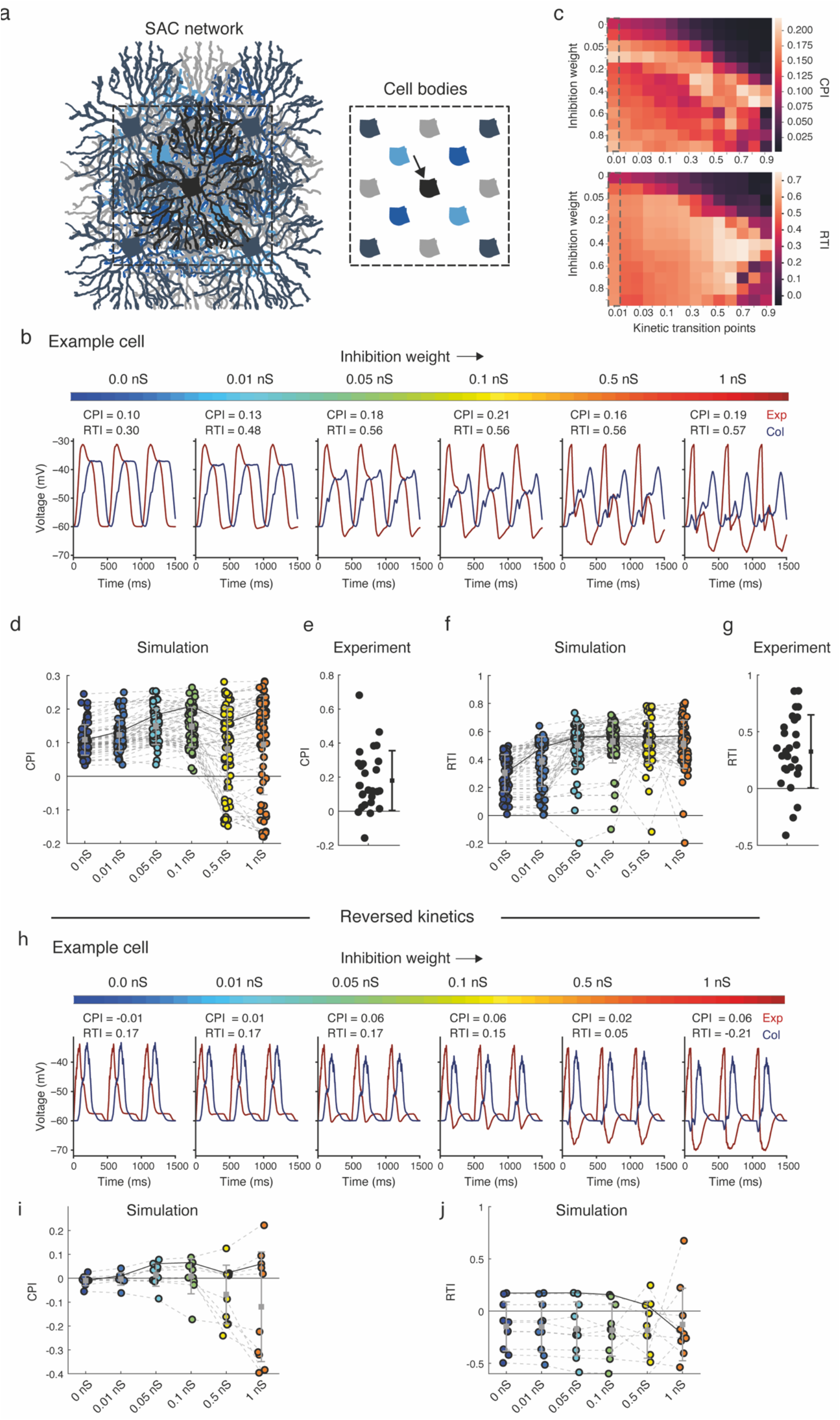
The strength of lateral inhibition differentially affects the amplitude and rise time of SAC responses. **a.** Simulated SAC network, consisting of two layers of replicated SAC sheets (scales of grey and blue). Right: illustration of all cell bodies plotted for clarification. The recorded simulated SAC (black, indicated by an arrow) resides in the center of the network. **b**. The voltage response of an example simulated central SAC (same as simulated SAC 1 from **Figure 3**) to expanding (red) and collapsing (blue) rings, shown for six different inhibition weights (colormap on top). **c.** Matrix of color-coded CPI (top) and RTI (bottom) values as a function of the different inhibition weights and kinetic transition points. The grey dashed box indicates the kinetic transition points used in (**b)**. **d**. The CPI of a population of simulated SACs (n=76) embedded in SAC networks with different inhibition weights. **e.** The CPI of the experimentally recorded SAC population (n=27 cells). **f**. Same as **d** but for RTI values. **g**. Same as **e** but for RTI values. See also **Supplementary Figure 7**. **h-j**. As in **b, d, f** but for simulated SACs with reversed spatiotemporal excitation: transient in proximal and sustained in distal processes.

We then used the 76 simulated SAC networks to investigate how the strength of lateral inhibition in these networks affects SAC CF preference in terms of both amplitude and kinetics. We compared the CF preference of the SACs while embedded across six network configurations: in the absence of lateral inhibition and when inhibition gradually strengthened to 1 nS. The CPI values were positive even in the absence of functional inhibition from the network, reflecting the criteria for cell inclusion. Interestingly, the CPI values moderately increased with inhibition strength, but with stronger inhibition (>0.1 nS) the CPI values of many cells tended to decrease, and the variability within the population increased (**Figure 4d**). The CPI values extracted from the physiological data were often higher than those of the simulated SACs but were also more variable (**Figure 4e**). The response kinetics were more affected by lateral inhibition, showing a monotonic increase in RTI values as inhibition strength increased and reaching a plateau around 0.1 nS (**Figure 4f**). The RTI values derived from the experimental data were similar to the values received in the simulations when stronger inhibition was defined (**Figure 4g**, further comparisons of simulation and experimental data are shown in **Supplementary Figure 7**). Timing of inhibition from SAC to DSGC was shown to play an essential role in DSGCs’ directional response (Ankri et al. 2020; Hanson et al. 2019). Thus, by delaying the response of SACs to collapsing motion, SACs’ intra-inhibitory connections act as another mechanism supporting direction selectivity in DSGCs.

Can inhibition per se generate CF preference? We chose non-CF preferring SACs from the original pool of cells from any generation of the genetic algorithm run to answer this. None of these SACs (n=12 cells) displayed CF preference when lateral inhibition was added to the SACs network, regardless of inhibition strength, and a portion of the cells even displayed negative CPI and RTI values (**Supplementary Figure 8**). This was also true for SACs depicted from the reversed kinetic simulation (with transient proximal and sustained distal inputs, see **Figure 4h-j;** n=10 cells). Thus, our simulation implies that while inhibition enhances SAC CF response, it is insufficient to generate it.

### Simulating the direction selective circuit: the role of SAC-DSGC and SAC-SAC inhibitory connections in DSGC direction selectivity

So far, we have investigated the mechanisms that underlie CF preference in SAC processes. It was recently suggested that SAC intra-inhibitory connections are more influential on the DSGC directional tuning than on the SAC response amplitude (Chen et al. 2020). We therefore shifted our focus to DSGCs, which directional response is thought to rely both on SAC CF preference and on the asymmetric wiring from SACs to DSGCs (**Figure 1a-c**; Wei 2018; Vaney et al. 2012; Mauss et al. 2017; Borst and Euler 2011; Briggman et al. 2011). Our goal was to dissect the contribution of each circuit component to DSGC’s direction selectivity. For this purpose, we used a reconstructed DSGC (Neuromorpho.org; ID: NMO_05318) and embedded it in a SAC network (see Methods for further details). In all conditions described below, the SAC-SAC inhibitory connections were either set to 0 nS (no intra-inhibitory connections) or to 0.1 nS (optimal strength as assessed by CPI and RTI; **Figure 4**). The SAC-DSGC inhibitory synaptic strength was fixed on 0.5 nS based on experimental data (Wei et al. 2011; see Methods). To determine the threshold for spiking, we conducted intracellular current-clamp recordings from DSGCs. Based on these recordings, we set the threshold for activation to 11 mV above baseline (**Supplementary Figure 9**).

We started by a network of CF-preferring SACs that contained intra-inhibitory connections and were randomly connected to the DSGC. With this random connectivity scheme, the circuit did not produce direction selectivity in DSGC responses, as assessed by bars moving in two opposite directions (**Figure 5a**). Next, we implemented the known asymmetric SAC-DSGC connectivity rule, with SAC processes preferentially connecting to DSGCs with a preferred direction antiparallel to the SAC process (Briggman et al. 2011). To implement this rule, we set the probability function of synapse formation between SAC and DSGC as the inverse cosine of the similarity between the direction of the SAC process (relative to the soma) and the preferred direction of the DSGC (**Figure 5b**). When running the simulation in the absence of SAC-SAC inhibition, the DSGC displayed direction selectivity, as preferred direction motion evoked stronger depolarization in the DSGC than null direction motion (**Figure 5c,** left). The inclusion of SAC-SAC inhibitory connections increased the response in the preferred direction (referred to here as PD activation). Still, it did not change the tuning strength measured by the direction selective index (DSI, see Methods) (**Figure 5c,** right and **5e**).

**Figure 5.**
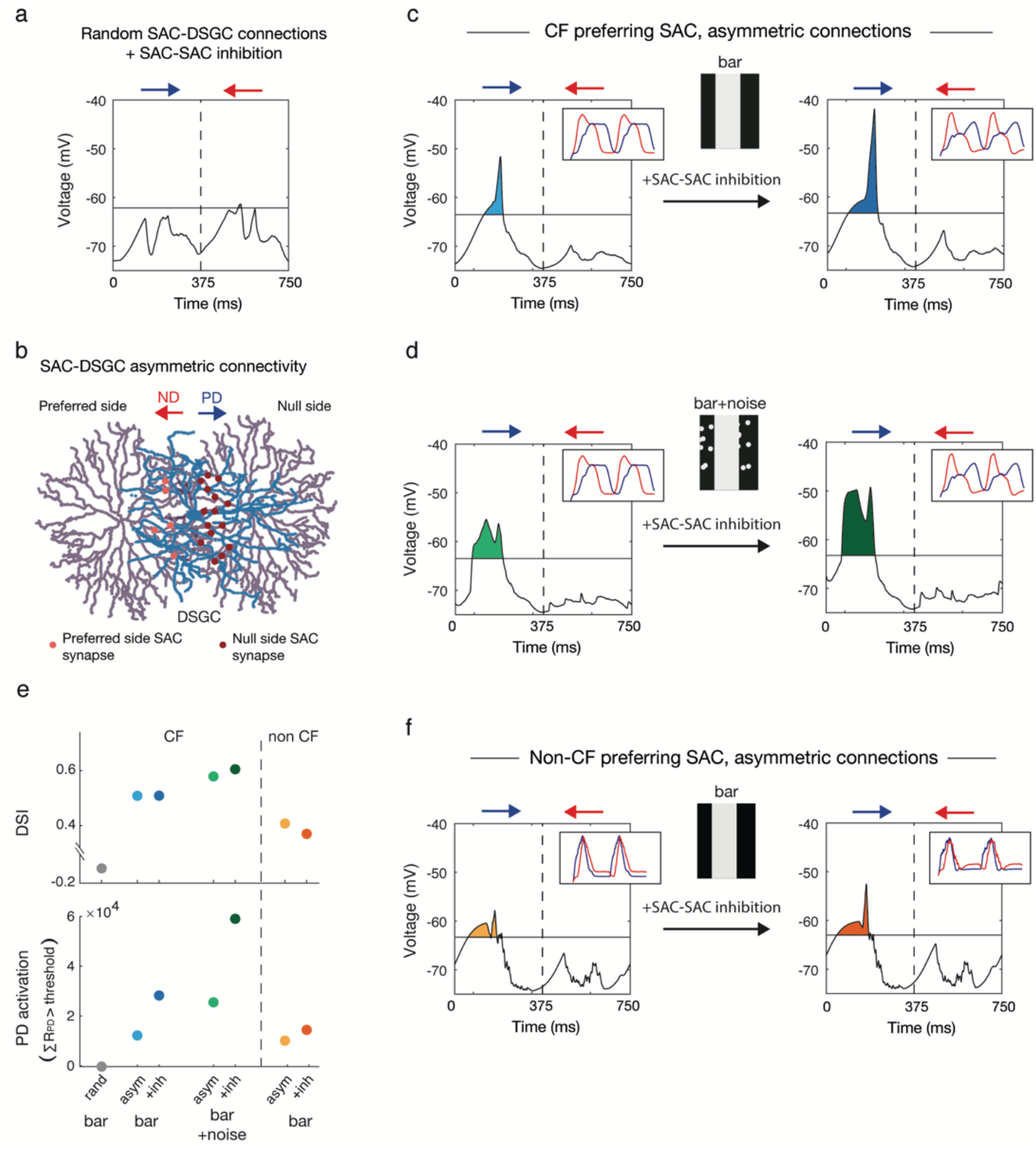
Asymmetric SAC-DSGC connections, but not CF SAC preference, are required for DSGC’s direction selectivity. **a.** A DSGC was randomly connected to a network of CF-preferring SACs (**Figure 4a**) and presented with a bar moving in the preferred (PD, blue) and null directions (ND, red). Voltage traces represent the DSGC’s somatic voltage. **b.** Schematic of two SACs (purple) forming asymmetric GABAergic synapses (red) onto a DSGC (blue). SAC-DSGC synapses were defined with higher probability when the SAC’s process and the DSGC’s PD were antiparallel. **c.** The response of the simulated DSGC when connected to the SAC network following the asymmetric connectivity rule, in the absence (left) and presence (right) of SAC-SAC inhibition. Insets represent the SAC waveforms to expanding and collapsing rings in each condition. **d.** Same as **c** but when random noise was added to the background of the visual stimulus. **e.** The direction selective index (DSI) and PD activation calculated from the above simulations. PD activation is calculated as the area above threshold during preferred direction motion. **f.** Same as **c** but for a non-CF SAC network.

SAC-SAC inhibitory connections were previously shown to play a unique role in direction selectivity in the presence of a noisy background (Chen et al. 2016; Chen et al. 2020). We relied on this knowledge to further examine the biological plausibility of our model. Random flickering spots were added to the background of the moving bar (See Methods), and the simulation was run on the network in the absence and presence of SAC-SAC inhibition (**Figure 5d**). The addition of the noisy background did not deteriorate the directional preference, and the DSGC maintained its preferred direction. Notably, while SAC-SAC inhibitory connections somewhat improved direction selectivity in the noiseless background, their effect was stronger in the presence of noise, in line with Chen and colleagues. This effect was most dominant for the PD activation (**Figure 5e**).

A recent study combined genetic manipulations with optogenetics to abolish SAC CF preference and demonstrated that direction selectivity in DSGCs is preserved (Hanson et al. 2019). RSME provided us with a unique opportunity to investigate the contribution of SAC CF preference to direction selectivity. For this, we generated a network of SACs from a non-CF preferring SAC (**Supplementary Figure 8a**). In accordance with Hanson’s study, the direction selectivity in DSGC was overall maintained, although to a lesser degree (**Figure 5e, f**). This result held both in the absence and presence of SAC-SAC inhibition, as the latter failed to generate CF preference in non-CF preferring SACs (**Figure 5f**, inset; **Supplementary Figure 8**).

Thus, DSGC’s response was stronger in the preferred direction in all simulations except for the random connectivity condition, as reflected by the positive DSIs (**Figure 5e**). This hints that null-side connectivity of SAC-DSGC is by itself sufficient to determine the preferred direction of the DSGC. SAC-SAC inhibition more strongly improved the directionality of the response when the bar was presented on top of a noisy background and had little or no effect when the bar was presented on a noiseless background or when the SAC was not CF preferring to begin with.

## Discussion

Although our understanding of the retinal connectivity pattern is rather solid (Helmstaedter et al. 2013; Bae et al. 2018; Briggman et al. 2011) and an abundance of physiological data have been collected to decipher retinal function, many of the mechanisms that underlie retinal computations are yet to be deciphered. Pure experimental approaches are imperative but often insufficient to reveal the contributions of neurons’ biophysical properties, input dynamics, and network connections to their functions. Therefore, the utilization of computational tools in retina research is essential. The dynamic nature of retinal functions (Rivlin-Etzion et al. 2018; Wienbar and Schwartz 2018) and the topographic variations in visual processing (Heukamp et al. 2020; Baden et al. 2020) further emphasize the need for robust simulation tools to overcome uncontrolled experimental caveats. While most modeling frameworks balance the biological details with the model’s complexity, RSME was designed to provide a modeling environment for retinal circuitry, combining a high level of biological details and support for large-scale retinal circuits. RSME allows building any neuronal circuit with detailed morphology, biophysical properties, and connectivity rules. The user can generate visual patterns, stimulate the retinal circuit and acquire the voltage in each of the neurons’ compartments. RSME can be extended to model other, non-retinal neuronal circuits, and its visual stimulation module may be used to generate patterns of optogenetic activation. Here, we exploited the exploratory power of RSME to decipher the underlying mechanisms of retinal direction selectivity and the origin of SAC CF preference.

SAC CF preference is a key component in the DSGC computation and various hypotheses, from intrinsic properties to network mechanisms, have been raised over the years to explain its source. Using RSME, we were able to dissect the network mechanisms, which rely on the architecture of SAC excitatory and inhibitory inputs. In the mouse retina, excitatory inputs from bipolar cells to SAC are confined to the proximal parts of its processes (Ding et al. 2016; Vlasits et al. 2016; Ankri et al. 2020). In addition, there is evidence that excitation to SACs is spatiotemporally diverse, with sustained bipolar cells preferentially innervating SAC proximal processes and transient bipolar cells innervating processes further from the cell soma (Greene et al. 2016; Kim et al. 2014; Fransen and Borghuis 2017; Ankri et al. 2020), although this remains controversial (Stincic et al. 2016; Ding et al. 2016). Using a genetic algorithm, we demonstrated that this spatiotemporal arrangement of the excitatory inputs is sufficient to evoke SAC CF preference. Notably, when forcing the reversed arrangement, where transient inputs innervate the proximal processes and sustained inputs innervate processes further from the cell soma, the algorithm did not converge and no CF preferring SAC was found. This result was obtained although we kept the anatomical constrain of denser inputs in the proximal processes, emphasizing the role of precise input kinetics in mediating SAC CF response.

Inhibitory GABAergic connections between neighboring SACs have been suggested to contribute to their CF preference by reducing SAC responses to centripetal motion (Lee and Zhou 2006; Poleg-Polsky et al. 2018), but opposing evidence was also found (Hausselt et al. 2007; Oesch and Taylor 2010; Chen et al. 2016; Hanson et al. 2019). We revealed that lateral inhibition strengthened SAC CF preference but differentially controlled different aspects of it. On the one hand, weak inhibition enhances asymmetries in SAC response amplitudes, and too strong inhibition can impair this asymmetry. On the other hand, kinetic differences monotonically increase with inhibition strength (**Figure 4**). This suggests that the inhibitory connections between neighboring SACs should be carefully tuned to optimally recruit both mechanisms for direction selectivity – SAC inhibition strength and timing.

We tested RSME by recapitulating experimental results showing that inhibitory connections within the SAC network are required to optimize DSGC’s direction selectivity on a noisy background (Chen et al. 2016; Chen et al. 2020). The rationale is that the addition of random flickering dots to the background of the moving bars effectively activates the SACs in the network such that the inhibitory connections between them are strongly engaged. Chen et al. demonstrated that the SAC-DSGC inhibitory synapse undergoes a short-term depression, which is more likely to happen in a noisy background. They suggested that the SAC-SAC inhibition prevents this depression, maintaining null-motion inhibition to DSGC and thereby maintaining direction selectivity (Chen et al. 2020).

We also exploited RSME to test whether a network of SACs that show no CF preference can induce direction selectivity in DSGC. Previously, selective reduction of SAC CF preference via elimination of SACs’ GABA-A receptors was found to decrease direction selectivity in DSGC (Chen et al. 2016). Yet, another study that used the same elimination model with SACs optogenetic activation showed that although inhibitory inputs to DSGC were symmetric (indicating loss of SAC CF preference), direction selectivity in DSGC was still maintained (Hanson et al. 2019). Our simulation results revealed direction selectivity in DSGC even when its innervating SACs lacked any directional preference, although this selectivity was reduced compared with the network of CF preferring SACs. Considering the positive effects of SAC-SAC inhibition on SAC CF preference and on DSGC’s direction selectivity provides a direct link between the two. Notably, our results demonstrate that asymmetric SAC-DSGC connections are essential for generating direction selectivity, as no direction selectivity emerged when SAC-DSGC connections were randomly distributed.

The extensibility of RSME enables us to further investigate the classic direction selective circuit at various mechanistic levels and thereby tackle many of the open questions in the field. First, the intrinsic properties of the SAC, such as differential distribution of ion channels along its processes (Hausselt et al. 2007; Oesch and Taylor 2010; Ozaita et al. 2004), somatic activation of mGluR receptors (Koren et al. 2017), and the location-dependent expression of chloride transporters (Gavrikov et al. 2006), have all been suggested to contribute to SAC CF preference. RSME uses the benefits of NEURON in the implementation of detailed biophysical properties. Each of the abovementioned intrinsic properties can be applied to the simulation, allowing the user to differentiate the role of each ion channel and transporter without compromising on the details of the whole network. Second, SAC network density was shown to be fundamental for generating direction selectivity in DSGCs (Morrie and Feller 2017). RSME can control the SAC coverage factor, thereby exploring the relationship between DSGC tuning strength and SAC network density. Third, SACs release both GABA and acetylcholine. While SAC GABAergic input to DSGC is asymmetric, the cholinergic input to DSGC was shown to be symmetric and its role in mediating direction selectivity remains controversial (Lee et al. 2010; Yonehara et al. 2011; Vaney et al. 2012; Pei et al. 2015). Different studies have hypothesized that acetylcholine specializes in maintaining direction selectivity for specific stimuli, depending on contrast level, stimulus size, background, or spatial frequency (Sethuramanujam et al. 2018; Grzywacz et al. 1998; Lee et al. 2010; Poleg-Polsky and Diamond 2016a; Chen et al. 2016). RSME allows the implementation of acetylcholine as additional output from SACs, independent from its GABAergic output. Its visual stimulation module can generate and explore a battery of stimuli and determine the contribution of the cholinergic signal to direction selectivity in different conditions. Forth, it was shown that direction selectivity is maintained over a wide input range, such as various speeds or light intensities (Chen and Wei 2018; Chen et al. 2020; Yao et al. 2018). RSME visual stimulation module can be further used to vary the characteristics of the moving stimuli to dissect the set of mechanisms that enable a stable directional response. Finally, the segregation to On and Off pathways is fundamental to the retinal structure. Here, we ignored the Off layer and investigated direction selectivity in the On pathway under the assumption that On and Off inputs are independent and mirror-symmetric. Yet, accumulating evidence hints towards discrepancies in the computations of the On and Off pathways (Wei 2018), and there is specific evidence that lateral inhibition between SACs, as well as inhibition arising from wide-field amacrine cells, differentially affect On and Off direction selective computations (Chen et al. 2016). RSME supports the implementation of another layer activated by light decrements, allowing testing the consequences of routing signals from both layers onto a single DSGC.

RSME provides a versatile, accessible framework dedicated to retinal research. Previous models of the retina used various levels of abstraction. Theoretical models have helped explain the retina’s neuronal diversity (Ocko et al. 2018; Kastner et al. 2015). Linear-nonlinear (LN) models have successfully reproduced the firing rates of retinal ganglion cells (Chichilnisky 2001; Carandini et al. 2005). Generalized integrate-and-fire (gIF) models have captured the influence of spike history and accounted for variability in retinal responses (Pillow et al. 2005). Realistic compartmental models of individual neurons have elucidated the contribution of ion-channels and morphological properties to retinal computations (Poleg-Polsky and Diamond 2016b; Ding et al. 2016). Each of these approaches advances our understanding of the system. Still, end-to-end dissection of retinal functions and their underlying mechanisms requires multiscale models: combining circuit connectivity and synaptic dynamics with realistic morphology and biophysics. Although various modeling frameworks have attempted to simulate such models (Martínez-Cañada et al. 2016; Bálya et al. 2002; Smith 1992; Wohrer and Kornprobst 2009), it has been challenging to study the biophysical mechanisms of retinal function in the context of circuit-level interactions. To the best of our knowledge, RSME is the first versatile framework that incorporates retinal neural networks with realistic synaptic connections, detailed morphological, biophysical and topological constraints of each neuron, as well as a wide range of visual inputs. Together, these make RSME an attractive tool for investigating the abundance computations the retina performs, from light adaptation via direction selectivity to approach sensitivity and motion prediction (Gollisch and Meister 2010) and evaluate their underlying mechanisms. RSME is available online at https://github.com/NBELab/RSME.

## Materials and Methods

### Animals

Trhr-EGFP mice (http://www.mmrrc.org/strains/30036/030036.html) (Rivlin-Etzion et al. 2011) and mGluR2-EGFP mice (Watanabe et al. 1998) were used for recordings. Mice were from either sex and various ages, from 4 weeks to 12 months. All procedures were approved by the Institutional Animal Care and Use Committee (IACUC) at the Weizmann Institute of Science.

### Electrophysiological recordings

Dark-adapted mice were anesthetized with isoflurane and decapitated. The retina was extracted and dissected in oxygenated Ames medium (Sigma, St. Louis, MO, USA) under dim red and infrared light. The isolated retina (dorsal part) was then mounted on a 0.22 mm membrane filter (Millipore) with a pre-cut window to allow light to reach the retina and put under a two-photon microscope (Bruker, Billerica, MA, USA) equipped with a Mai-Tai laser (Spectra-physics, Santa Clara, CA, USA) as previously described (Warwick et al. 2018; Ankri et al. 2020). For SAC recordings, we used mGluR2-EGFP mice (Watanabe et al. 1998) and for DSGC recordings, we used TRHR-EGFP mice (Rivlin-Etzion et al. 2011), which express GFP in SACs and posterior preferring DSGCs, respectively. GFP-expressing cells were targeted for recordings with the laser wavelength set to 920 nm to minimally activate photoreceptors using a 60x water-immersion objective (Olympus, Tokyo, Japan). The isolated retina was perfused with warmed Ames solution (32–34 °C) and equilibrated with carbogen (95% O2:5% CO2). Current-clamp recordings from both SACs and DSGCs were made using 5–9-MΩ glass pipettes containing (in mM): 110 KCl, 2 NaOH, 2 MgCl2, 0.5 CaCl, 5 EGTA, 10 HEPES, 2 ATP, 0.5 GTP and 2 Ascorbate (pH = 7.2 with KOH; Osmolarity = 280). Extracellular spike recordings from DSGCs were made in loose cell-attached mode using a pipette filled with Ames solution. Data were acquired using pCLAMP10, filtered at 2 kHz and digitized at 10 kHz with a MultiClamp 700B amplifier (Molecular Devices, CA, USA) and a Digidata 1550 digitizer (Molecular Devices). The electrophysiological data of SAC recordings presented here combines published and new data (Ankri et al. 2020). All DSGC recordings are new data.

### Experimental Light Stimuli

Visual stimuli were generated using MATLAB and the Psychophysics Toolbox. Stimuli were projected to the retina by a monochromatic organic light-emitting display (OLED-XL, 800 x 600 pixels, 85 Hz refresh rate, eMagin, Bellevue, WA, USA) as previously described (Ankri et al. 2020). SAC CF preference was assessed by the presentation of expanding and collapsing rings centered on the SAC soma, projected via a 20x objective for 5 sec and repeated five times in a pseudo-random order. The spatial frequency of the rings was 450 μm/cycle and the temporal frequency was 2 Hz, resulting in 900 μm/sec velocity. The SAC soma was masked by a 25 μm radius grey spot (Euler et al. 2002) and the first cycle was removed from the analysis. DSGC directional tuning was assessed by its responses to bars moving in the preferred and null directions (900 μm/sec; 300 μm width; 1200 μm length) repeated four times in a pseudo-random order. DSGCs’ spike properties were extracted from their response to a 2-sec static bright spot (100 μm diameter) centered on the cell soma and projected through a 60x objective.

### Data analysis

SAC tuning was assessed by their CF preference index (CPI):

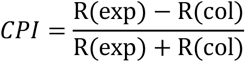

where R(exp) and R(col) are the amplitudes of the response to expanding and collapsing rings, respectively. SAC response kinetics were assessed using the response rise time, measured as the delay between the initial response (20% of the peak) and the maximum point (Ankri et al. 2020). The rise times were then used to calculate the rise-time index (RTI) as a measure of directional asymmetries in SAC response kinetics:

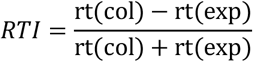

where rt(col) and rt(exp) are the rise times calculated in response to collapsing and expanding rings, respectively. Similar measurements were used for experimental and simulation data. To calculate the parameters of DSGC’s spiking, we measured the baseline voltage of the cells as the mean voltage 0.5 sec before spot illumination following the removal of fast-spiking events. Spikes evoked during the 2 seconds of light presentation were detected and removed, and the remaining voltage trace was filtered (Savitzky-Golay filter, 3^rd^ order). Traces that showed a baseline >-45 mV were removed. The spiking threshold was calculated as the maximum depolarization on which the evoked spikes were riding. 151 spikes were detected for this analysis, from 15/16 intracellularly recorded DSGCs: one cell was removed from the study due to low spiking amplitudes (<50 mV).

For simulated DSGCs, direction selectivity was assessed by two parameters: direction selective index (DSI) and PD activation. DSI was evaluated by:

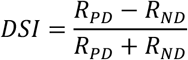

where *R_PD_* and *R_ND_* are the areas under the voltage trace in the preferred and null direction, respectively. PD activation was measured as the area under the voltage trace and above the spiking threshold in the preferred direction. The threshold was set to 11 mV above minimum voltage to fit with our experimental measurements.

The sustained transient index (STI) was calculated for each synapse by simulating its dynamics based on release probability and refilling rate for 250 ms. Since the synapses are stochastic, we simulated each synapse 50 times and averaged the peristimulus time histogram (PSTH) (with 10 ms bins). Using the PSTH, we calculate the sustained transient index:

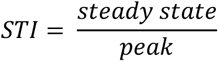

where the peak was set to the value in the first bin after the simulation starts, and the steadystate was assigned to the value in the last PSTH’s bin.

We made a 2d interpolation table for the sustained transient index over the entire range of release probabilities (1.56 × 10^-4^, 0.25) and refilling rates (5.12 × 10^-5^, 1.0) with 51 equally logarithmically spaced points along each axis. We estimated the index for each synapse by first calculating its release probability and refilling rate (both depend on distance from the soma) and then using linear interpolation over the table.

### RSME framework

RSME encapsulates NEURON (Carnevale and Hines 2006) and its XML-based specification interface was inspired by NeuroML and used SBML (Systems Biology Markup Language) for the description of mathematical models (Gleeson et al. 2010; Hucka et al. 2003). RSME initialization follows a top-down approach and is defined in 5 layers:

I. **Meta parameters**. Simulation unique identifier, simulation duration, and XML file paths for layers II-V. Model parametrization is based on a series of four XML files, providing input for stimulation patterns as well as architectural, morphological, and biophysical specifications (**Supplementary Figure 1**). The simulation duration used in this work was 1,500 ms.
II. **Visual stimulation**. Stimulation parameters include stimulation type (e.g., moving bars or circular moving rings), stimulation field (size and location), spatial and temporal frequencies (when applicable) and duration. The stimulation pattern is parsed and conveyed into a gray-scale array, specifying the illumination level for each point in space and time (**Supplementary Figure 1**, right). The visual stimulation is transferred to the cells via the light- and dark-activated synapses (see Biophysical properties).
III. **Network architecture**. Inspired by NeuroML, RSME uses *populations* of identical neurons, assigned with cell ID (defined in layer IV) and spatial arrangement. An arrangement can be specified by indicating a location in 3D space for each cell or using a template for an entire cell population. For example, cells can be arranged in a grid or layers of grids, defined by grid location, the number of cells and spacing. RSME supports multiple populations in a single run, and connectivity rules can be defined within a population of neurons (termed *intra-synapses)* or between populations of neurons (termed *projections).* Synaptic connections obey the connectivity rules and are restricted to intersections between the pre- and post-synaptic cell. Intersections are identified based on spatial proximity within a threshold value. Note that a simulation may contain all types of retinal neurons, starting with photoreceptors and ending with retinal ganglion cells. However, depending on the investigated cells and circuits, one may omit the populations of photoreceptors and even bipolar cells to speed up simulation run-time. In this case, synaptic inputs to the simulated cells should be designed to mimic input from bipolar cells (**Supplementary Figure 1**). The simulations described in this work used this option.
IV. **Cell morphology**. Each cell’s morphology is assigned with a unique identifier (ID). Morphology for each cell can be either defined using a raw reconstruction file (asc, swc filetypes) or precompiled morphology reconstruction (NEURON Hoc file). Since cell reconstruction tools often have a limited resolution, RSME supports morphology ad hoc correction functions.
V. **Biophysical properties**. Each cell population is defined with its ion channels and cellular properties, including cytoplasmic resistivity, membrane capacitance and conductance dynamics of the various channels. Each channel can be distributed either uniformly or following a density function. All mathematical functions (e.g., density and distribution functions) are defined using the Systems Biology Markup Language (SBML), a widely accepted standard for bioinformatics (Hucka et al. 2003). For example, one can electronically isolate the soma with low potassium conductivity or define a high conductance of Nav1.8 sodium channels in distal dendrites (**Supplementary Figure 2b**). This layer is also used to specify synapse properties. RSME supports retina-tailored specifications such as light- and dark-activated synapses. Dark-activated synapses can be defined directly on photoreceptors, mimicking the photoreceptor’s dark current. When the simulation omits photoreceptors and bipolar cells, light- and dark-activated synapses can be defined to mimic the input coming from On and Off bipolar cells, respectively. In this case, the distribution of the light- and dark-activated synapses should follow the anatomical inputs from bipolar cells, and their activation is determined by the spatiotemporal pattern of the visual stimulation. Since different bipolar cells show different response kinetics (Franke et al. 2017), RSME enables the definition of each synapse’s transient/sustained properties, indicating the temporal dynamics of the response relative to stimulus presentation.

RSME implements a flexible visualization tool for model exploration. 2-dimensional morphological projections can be visualized for each layer in either full (all sections are shown) or soma mode (where only somas and bounding boxes are shown, **Supplementary Figure 3**). Cell morphologies with synapse locations and weight distribution can also be visualized so that each cell in the model can be visualized as a whole – from cytoplasmic resistance to channel distribution (**Supplementary Figure 2**). RSME can generate a series of images listing all defined properties for each cell with a single command line. See the project’s Github for further information.

### Single-cell SAC modeling

To assess the CF preference of simulated SACs, we generated a visual stimulation pattern of expanding and collapsing rings. The stimulation field was set to 315 x 315 μm, and the rings’ position was centered on the cell soma; in our simulation, this location was x = 135 μm and y = 125 μm. Stimulus frequency was set to 2 Hz. Visual stimulation was defined to operate via bipolar cell synapses. These functions, called *alternating_expanding_circles* and *alternating_collapsing_circles,* are provided in the *Stimuli_visual_pattern* library, which we designed as part of RSME. These visual patterns are defined using functional programming: the functions calculate the phase of the current stimuli and return another function, which given a phase and location, returns the appropriate illumination value. For expanding rings, the illumination phase is defined as (*t* – *delay*)*mod*(2 · *time_c_*) < *time_c_*, where *t* is the current simulation time, *delay* is the prespecified delay time for the stimulation (set to 0 in our simulation), *time_c_* is half the stimulation period, and *mod* is the modulus operator. The illumination phase is similarly defined for collapsing rings with the < operator changed to >. Within an illumination phase, we defined an effective circular field by scaling the stimulation field by 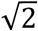 to account for the fact that a circle should expand far enough to cover a squared area of illumination. Other supported features are given in the project’s documentation file. Here, network specification included a single instance of a SAC, defined in the morphology specification file. We also defined an ad-hoc morphological correction function, which divided all sections’ radii by two to correct mislabeled morphology tracing.

SAC cytoplasmic specific resistance was set to 75 ohm*cm and specific capacitance was 1 μF/cm^2^. Section’s conductance was set to 0.00006 S/cm^2^ and resting potential to −60 mV. These parameters resulted in input resistance of 84 MΩ, within the experimental range observed in SAC neurons (**Supplementary Figure 4**).

Bipolar cell synapses onto the SAC neurons were modeled using a double exponent function with a rise and decay time of 0.89 ms and 1.84 ms, respectively (Singer et al. 2004) and reversal potential of 0 mV. The synaptic conductance of each synapse was set to determine the connectivity strength. The simulation we used included On SACs and their inputs from On bipolar cells.

During light offset, each synapse recruited 70 vesicles available for release (Singer and Diamond 2006). During illumination, vesicles were released at each time step from this pool with some probability (*p*), and the pool size decreased accordingly. In parallel, the pool was replenished with a refilling rate (*r*). Once the visual signal changed from light to dark, the readily releasable pool of each synapse was reset to 70. For each synapse, *p* and *r* were set by the following functions:

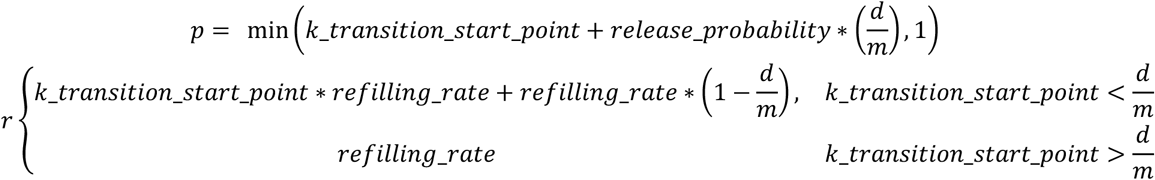

where *d* is the location of the synapse and *m* is the kinetic transition endpoint (*k_transition_end_point), refilling_rate* and *release_probability* are parameters that set the starting set point for *p* and *r*, and *k_transition_start_point* (kinetic transition start-point) dictates how *p* and *r* change with distance from the soma. The resulted kinetics were sustained-release around cell soma, and a gradual shift from sustained to transient at a distance *k_transition_start_point* from cell soma, which continued to change up to *k_transition_end_point,* from which the most transient kinetics is kept constant. For the reversed kinetics, we set *d* to *210-d.*

The biophysical specification included bipolar cell synapse distribution and passive parameters. The distribution of the bipolar cell synapses was defined (per μm) using the following rule:

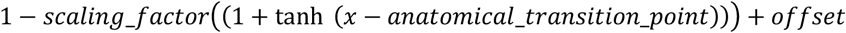

where *x* is the distance between a particular section and the soma. *scaling_factor, anatomical_transition_point* and *offset* are paramteres. The density at the soma (proximal density) is:

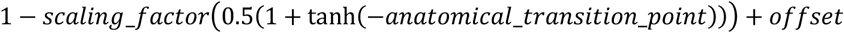

The density at the distal process was defined as:

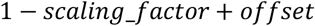

The values of these parameters were tuned using the genetic algorithm to maximize SAC CF preference in response to expanding and collapsing rings.

### Genetic optimization

We used a genetic algorithm to maximize SAC CF preference. We implemented the genetic algorithm using DEAP (Fortin et al. 2012) and explored an 8-dimensional operational space. Algorithm boundary parameters were:

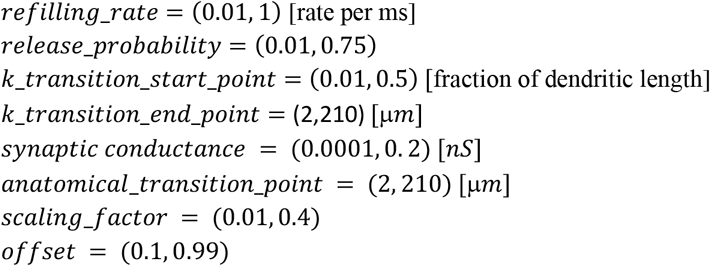

We defined three objectives for the optimization: (1) The difference between the maximal voltage that the SAC neuron reached during the centrifugal stimulus to that from the centripetal stimulus: Amp_CF_-Amp_CP_. (2) The difference between the rise time for the centripetal response and the centrifugal response (rise time defined as the time from 20% of the peak to the maximum point): RT_CP_-RT_CF_. (3) The difference between the maximal voltage for the centrifugal stimulus and −10 mV. If the maximal voltage was below −10 mV, this objective was set to zero. This criterion was meant to prevent non-physiological depolarizations above −20 mV. Additionally, in individuals where the difference between the maximal voltage for the centrifugal and the centripetal stimuli was lower than 0.04 mV, all objective values were set to −50. The weights of objectives 1-3 were 1, 0.3, and 0.08, respectively.

We started the genetic algorithm with a population of 100 individuals and ran it for 20-45 generations. The crossover and mutation probabilities were set to 0.4. For the crossover operation, we used mutFlipBit (Fortin et al. 2012). The mutation operation was applied by replacing the mutated variable with a variable sampled from a normal distribution with a mean and standard deviation equal to the mutated variable and half of the mutated variable, respectively. We used three different selection algorithms, Non-dominated Sorting Genetic Algorithm-II (Deb et al. 2002), Indicator-Based Evolutionary Algorithm (Van Geit et al. 2016), and one in which the individuals were ranked according to their total score. In one case, we run the algorithm with a fixed *k_transition_end_point*=135 μm. All three algorithms produced similar results.

### SAC network modeling

We tested the SAC network with expanding and collapsing rings. Both visual stimulation patterns were defined similarly to the single-cell simulations except for the stimulus’s center, which was set here to x = 260 μm and y = 250 μm to match the location of the central SAC in the network. The network was specified as two overlaying grids of cells. The first grid of cells was defined as a 3×3 2D grid (9 cells), with the bottom-left cell located at (0,0,0), and all other cells were organized with a 125 μm spacing between neighboring cell bodies horizontally and vertically. The second grid of cells was defined as a 2×2 2D grid (4 cells), where the bottomleft cell was cornered at (62,67,0) μm coordinate. Spacing between the cells was identical to the first grid. Overall, the network featured 13 SACs and the responses of the SAC in the center were analyzed and used in this study. Input distribution and kinetics were defined using the parameters we obtained from the genetic algorithm on the selected single SACs.

SAC-SAC GABAergic connections were modeled using a double exponent (NEURON’s Exp2Syn point process). Each synapse’s rise and decay times were 3 ms and 30 ms, respectively, and the reversal potential was −75 mV. When the voltage at the presynaptic section increased above −50 mV, the synapse was activated at 200 Hz.

### SAC-DSGC modeling

The SAC-DSGC model was defined similarly to the SAC network specified above, with the addition of a single DSGC. We only used the On layer to simplify the simulation. The DSGC was centered at (0,0,0) and defined with NEURON’s hoc file (available through Neuromorpho.org; ID: NMO_05318).

The visual stimulation in this case was a bar moving in the predefined preferred and null directions, with and without static noise in the background. Similar to the expanding and collapsing rings, the *alternating_bar* function returns a function that calculates the illumination level for each point in the network space at any given time. The bar’s length was 250 μm and its width was 600 μm; its motion velocity was set to 1000 μm/sec; and an x perimeter defined the area in which the bar is moving (here, 500 μm). The bar’s leading-edge location was calculated using *t* · *velocity* and the location of its trailing edge was calculated by adding the bar’s length to the leading-edge location. The bar was set to change its direction every (x perimeter + bar size)/bar velocity. To generate a visual stimulus where spots randomly flicker in the background of a moving bar, we defined a configurable mechanism for spots generation. Specifically, we regenerated 30 spots (25 μm in radius), randomly distributed across the field of view, in each flickering phase (15 Hz).

Note that the SACs and DSGC are not logically co-located in the same coordinate system. However, we can logically align them as we connect the two populations with synapses. Synapse formation was based on the x-y intersection (minimal distance of 5 μm) and the probability function of synapse formation between SAC and DSGC was set to the inverse cosine of the similarity between the direction of the SAC process (relative to the soma) and the preferred direction of the DSGC. The logical alignment was defined at x = 135 μm and y = 100 μm (so the DSGC’s soma was aligned to the soma of the central SAC). The conductance of SAC-DSGC synapses was estimated based on published data: dual recordings showed that SAC-DSGC connectivity strength is ~10 nS (Wei et al. 2011). Assuming ~20 contacts between a SAC and a DSGC, we set each synapse to 0.5 nS. Parameters for the SAC were chosen from the results of the genetic algorithm (see simulated SAC 1 in **Figure 3** and **Figure 4b**), defined with

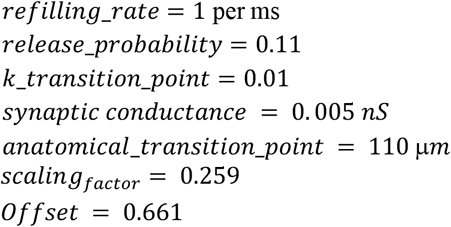

### Data processing and visualizations

We used MATLAB, python and NumPy (Harris et al. 2020) for data processing. Figures and neuron morphology visualizations were created using Matlab, Matplotlib (Hunter 2007), seaborn (Waskom et al. 2018), BlenderNEURON, and processed in Adobe Illustrator.

### Computer simulation

All simulations were run using NEURON 7.7 (Carnevale and Hines 2006). Genetic algorithm simulations were performed on the Blue Brain V supercomputer based on the HPE SGI 8600 platform hosted at the Swiss National Computing Center in Lugano, Switzerland. Each compute node was composed of an Intel Xeon 6140 CPU @2.3 GHz and 384 GB DRAM. All other simulations were performed on a 2.6 GHz 6-Core Intel Core i7, 16GB RAM, Mac computer.

## Acknowledgments

We thank the Blue Brain Project for computing time and Werner Van Geit for guidance in using the genetic algorithm. In addition, we thank members of the Rivlin lab for their comments and suggestions on the manuscript. This work was supported by research grants from the European Research Council (ERC starter No. 757732), the Israel Science Foundation (1396/15 and 2449/20), the Minerva Foundation, and by the Charles and David Wolfson Charitable Trust; Rolf Wiklund and Alice Wiklund Parkinson’s Disease Research Fund; Consolidated Anti-Aging Foundation; Dr. and Mrs. Alan Leshner; Dr. Daniel C. Andreae. L.A. is supported by ISEF, O.A. was supported by the ETH domain for the Blue Brain Project. M.R.-E. is incumbent of the Sara Lee Schupf Family Chair.

## Supplementary Figures

**Supplementary Figure 1.**
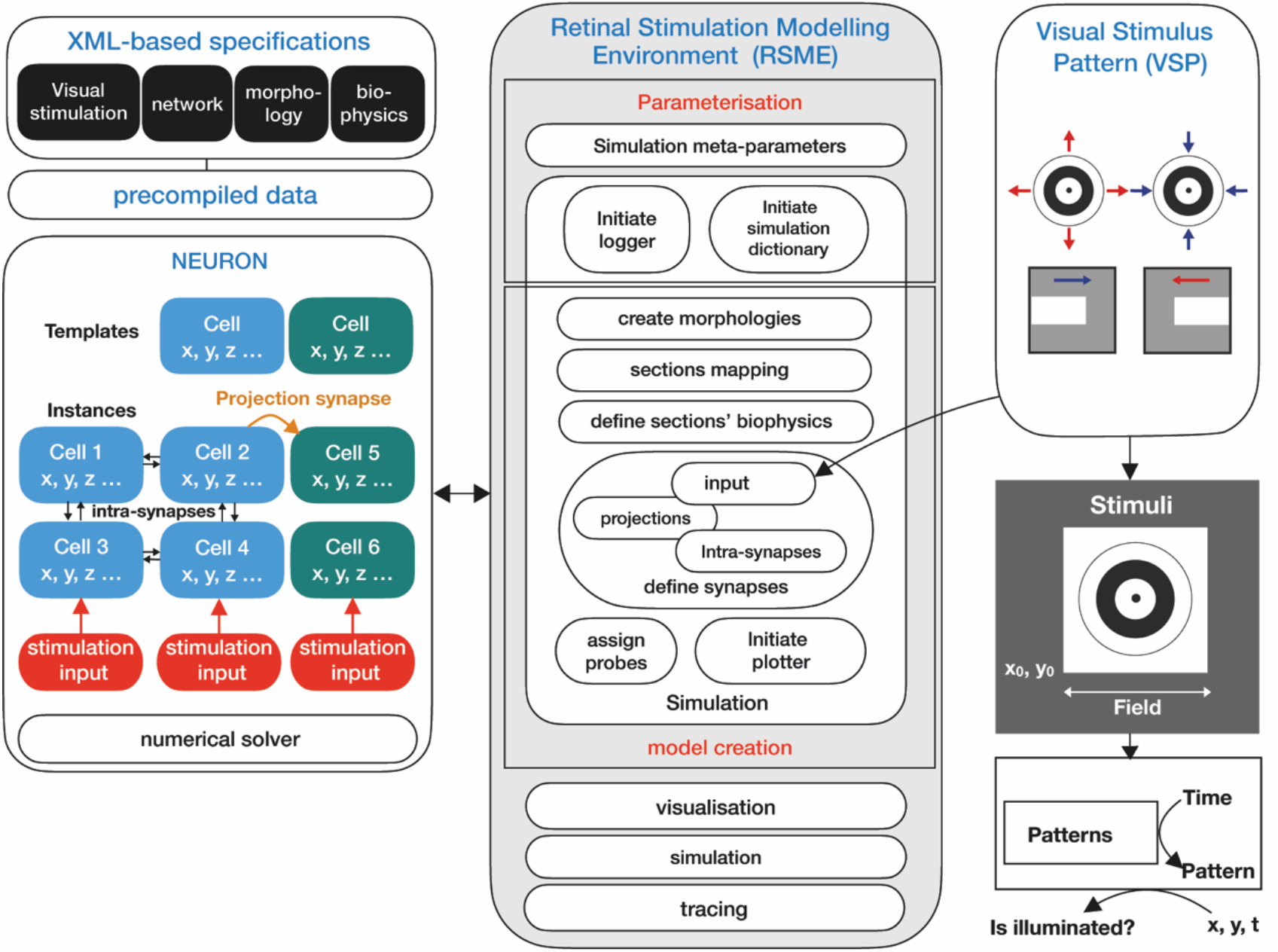
RSME top-down approach toward modeling. Modeling is initiated with XML-based specification and precompiled data. Parameters are parsed within RSME, which initializes a logger and supporting data entities and creates the model. Model creation is executed with NEURON and includes morphology creation and the definition of sections’ biophysics and synapses. RSME implements a visual stimulation pattern module, in which inputs such as expanding/collapsing rings and moving bars are provided to the network as structured light patterns. Model is simulated using NEURON’s solver, and the results are retrieved for analysis.

**Supplementary Figure 2.**
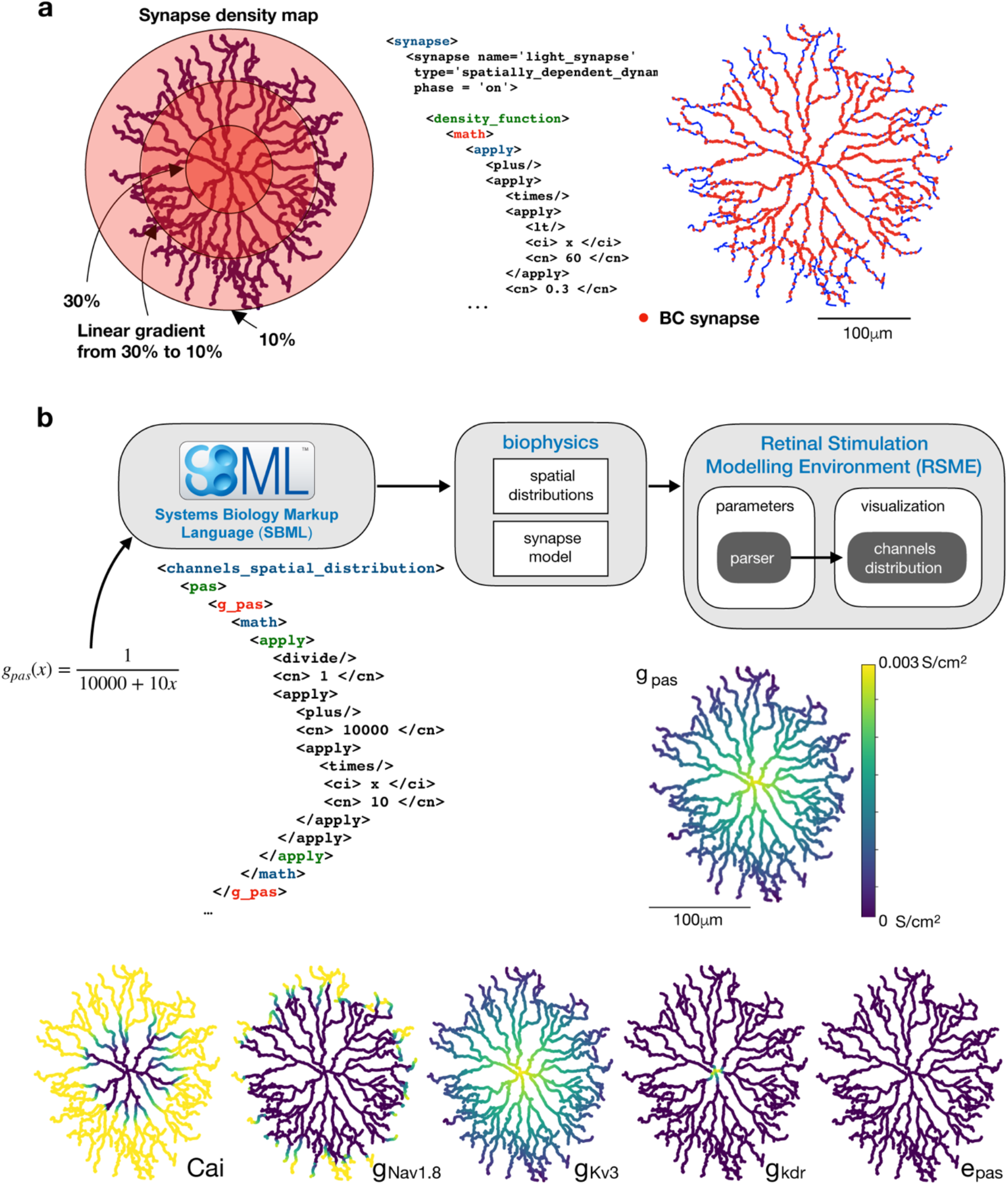
Synaptic input definition and biophysical specifications. **a**. An example synapse density pattern on a reconstructed SAC (Neuromorpho.org; ID: NMO_50993), where synapses are defined to 30% of the proximal sections (dark red shaded area), 10% of the distal section (light red shaded area), and linearly distributed in between. Synapse density is provided in an XML file, which is parsed and implemented by RSME. **b**. Example distributions of section’s passive conductance, reversal potential (g_pas_, e_pas_), and conductances of potassium, sodium and calcium channels (g_kdr_, g_kv3_, g_Nav1.8_, Cai). Values are according to SAC biophysical properties in (Ding et al. 2016). Parameter distributions are specified in XML using SBML and parsed within RSME, which provides distributed mapping of the parameters and visualization.

**Supplementary Figure 3.**
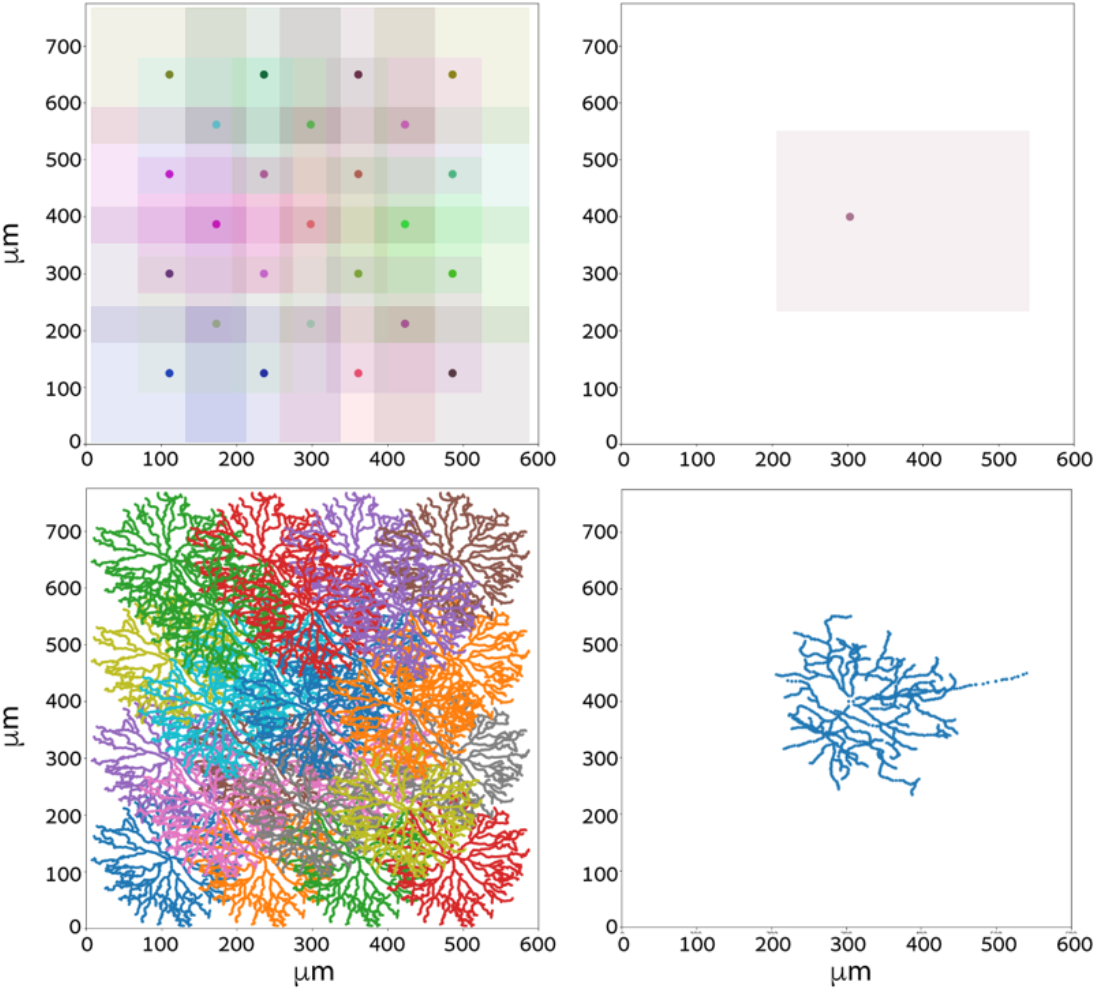
Cell morphology and arrangement. (Top) Soma plots for a SAC population comprised of two grids (containing 16 and 9 SACs) (left) and a single DSGC (right). Somata are indicated with dots and colored boxes indicate the size of the cell, demonstrating the overlapping regions between cells and the two grids arrangement of the cells. (Bottom) full morphological plots for the SAC network (left) and DSGC (right).

**Supplementary Figure 4.**
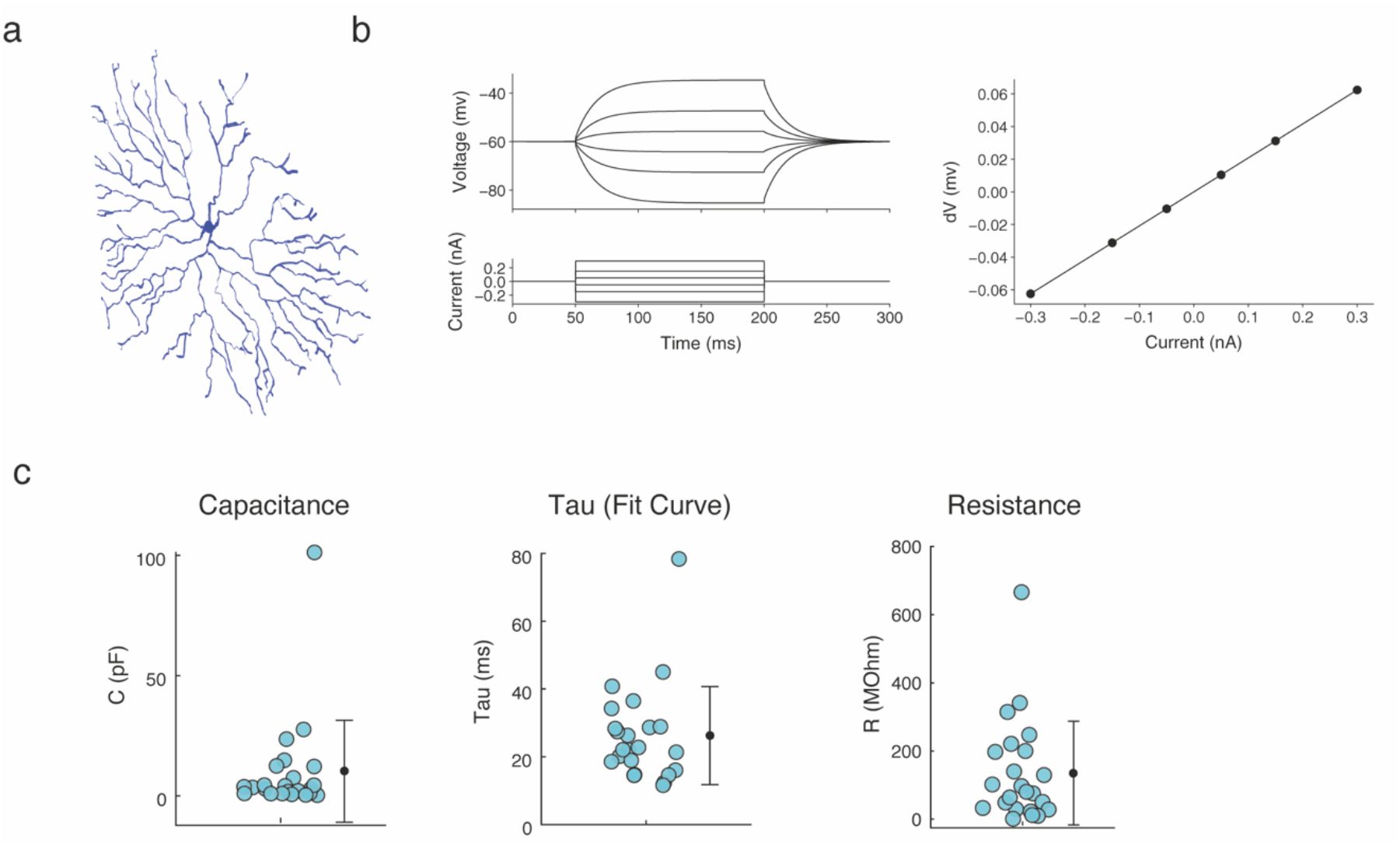
SAC passive properties. **a.** Model of a 3D reconstructed SAC (Neuromorpho.org; ID: NMO_139062). **b.** (Left) Different current injections and the corresponding voltage deflections for a simulated SAC which morphology is shown in **a**. (Right) The change in voltage as a function of the current that was injected to the simulated SAC; from the slope of this curve we can extract the input resistance of the cell (84 MΩ). **c.** The capacitance (left), time constant (middle) and resistance (right) of recorded SACs (N = 24). Values were extracted from the voltage response of the cells to a pulse of −5 pA during recordings.

**Supplementary Figure 5.**
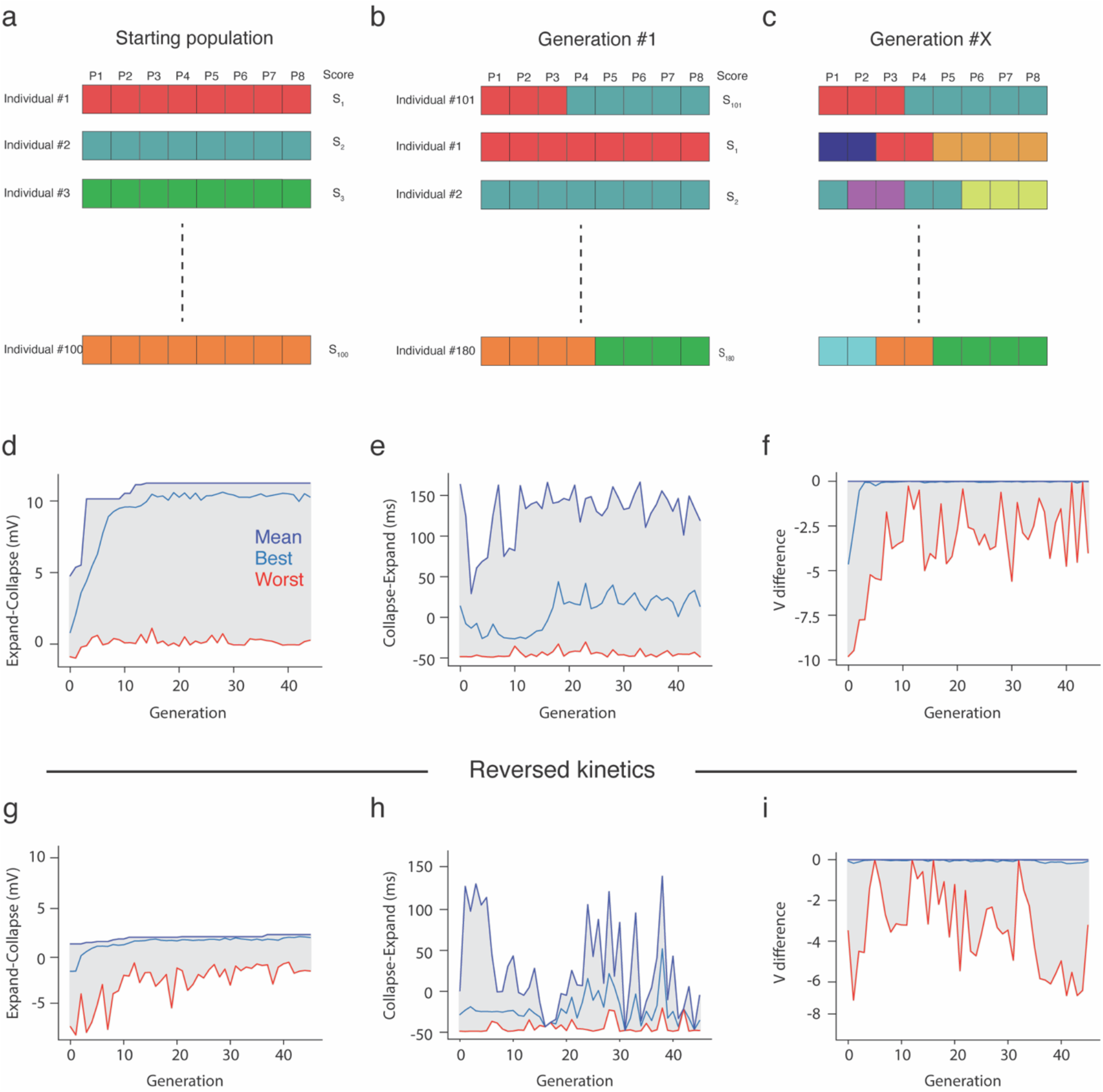
Genetic Algorithm parameter search. **a.** Initiation of the genetic algorithm population with 100 randomly selected individuals, each with eight parameters (see Methods). **b.** Illustration of the second generation. The best individuals from Gen 0 are copied to Gen 1 and the rest are a combination of different individuals from the previous generation (illustrated by individuals with combinations of colors). Random jitter (mutations) was added to some of the individuals. **c.** Same as in b for the X^th^ generation. **d-f.** Progression of different objectives, the response amplitude difference (**d**), rise time difference (**e**) and Voltage_score_ (**f**) (see Method), as a function of generation number. Higher values indicate a stronger centrifugal preference. **g-i.** Same as (**d-f**) but with the reversed input kinetics. Here, the genetic algorithm did not converge (as reflected by the rise time differences in **h**), and no CF preferring SACs were found (as reflected by the response amplitude difference in **g**).

**Supplementary Figure 6:**
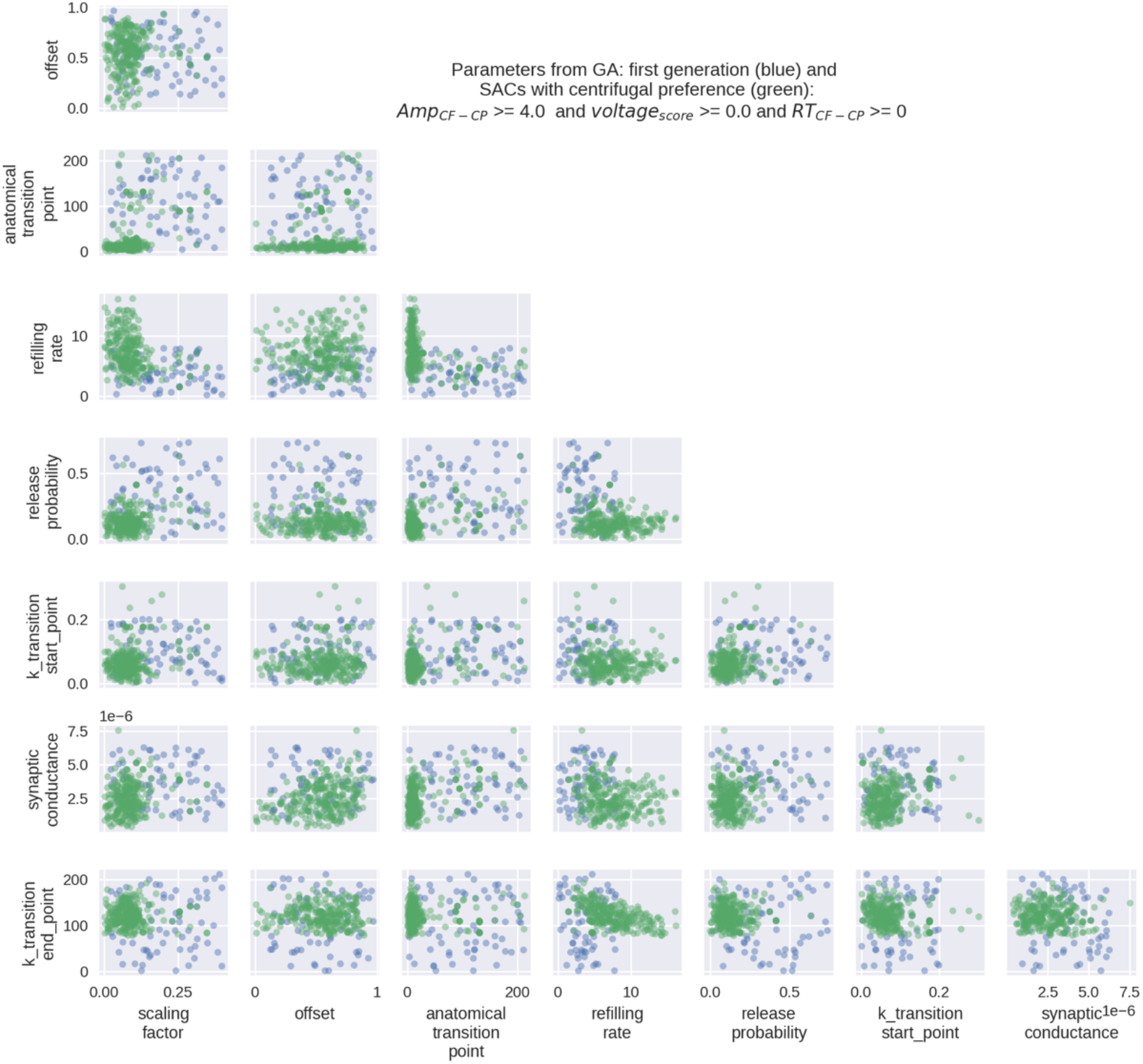
A scatter matrix of parameters from the genetic algorithm. The genetic algorithm was initiated to have a uniform distribution of parameters on all eight dimensions. The first generation shows the extent of these in blue, and in green are the parameters in the 1^st^ to 45^th^ generation, which resemble SACs in their input-output characteristics (see Methods). In the 45^th^ generation, the algorithm converges to several such realistic cells.

**Supplementary Figure 7.**
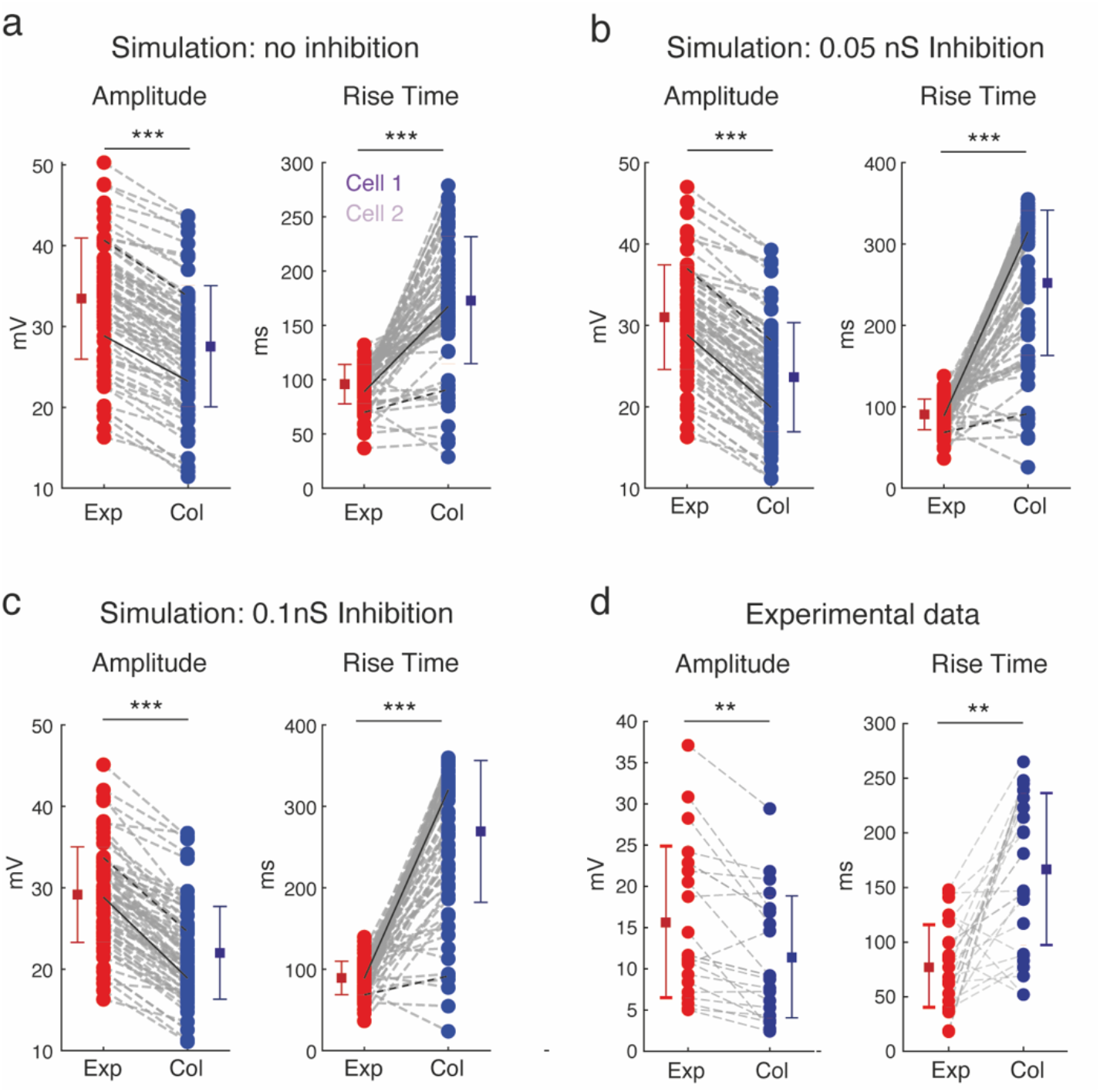
Comparison of the values extracted from simulated SAC responses and experimental data. **a.** The amplitude (left) and rise time (right) of the responses to expanding (red) and collapsing (blue) rings, when inhibition from the SAC network is set to 0 (Amplitude – expanding: 33.47±7.5 mV; collapsing: 27.56±7.5 mV; Rise time – expanding: 95.85±18.12 ms; collapsing: 173.3±58.54 mS). Dashed lines connect values from the same cell; solid and dashed black lines connect the values of example cells 1 and 2, respectively (**Figure 3**). Vertical bars denote the mean ± STD. **b**. Same as (**a**) but when 0.05 nS inhibitory conductance was implemented (Amplitude – expanding: 31± 6.42 mV; collapsing: 23.64±6.7 mV; Rise time – expanding: 90.69±18.9 ms; collapsing: 252.2±89.2 ms). **c**. same as (**b**) but for 0.1 nS inhibition (Amplitude – expanding: 29.16±5.86 mV, collapsing: 22±5.7 mV; Rise time – expanding: 89.18±20.52 ms; collapsing: 269.3±87 ms). **d**. same but for the population of experimentally recorded On-SACs (Amplitude – expanding: 15.3±8.8mV; collapsing: 11.3±7.4mV; Rise Time – expanding: 78.19±37.74; collapsing: 166.89±-69.52). Note that both amplitudes and rise times are in the range of the simulation results. (Statistical tests: student’s paired t-test, **p < 0.01; ***p < 0. 0005)

**Supplementary Figure 8.**
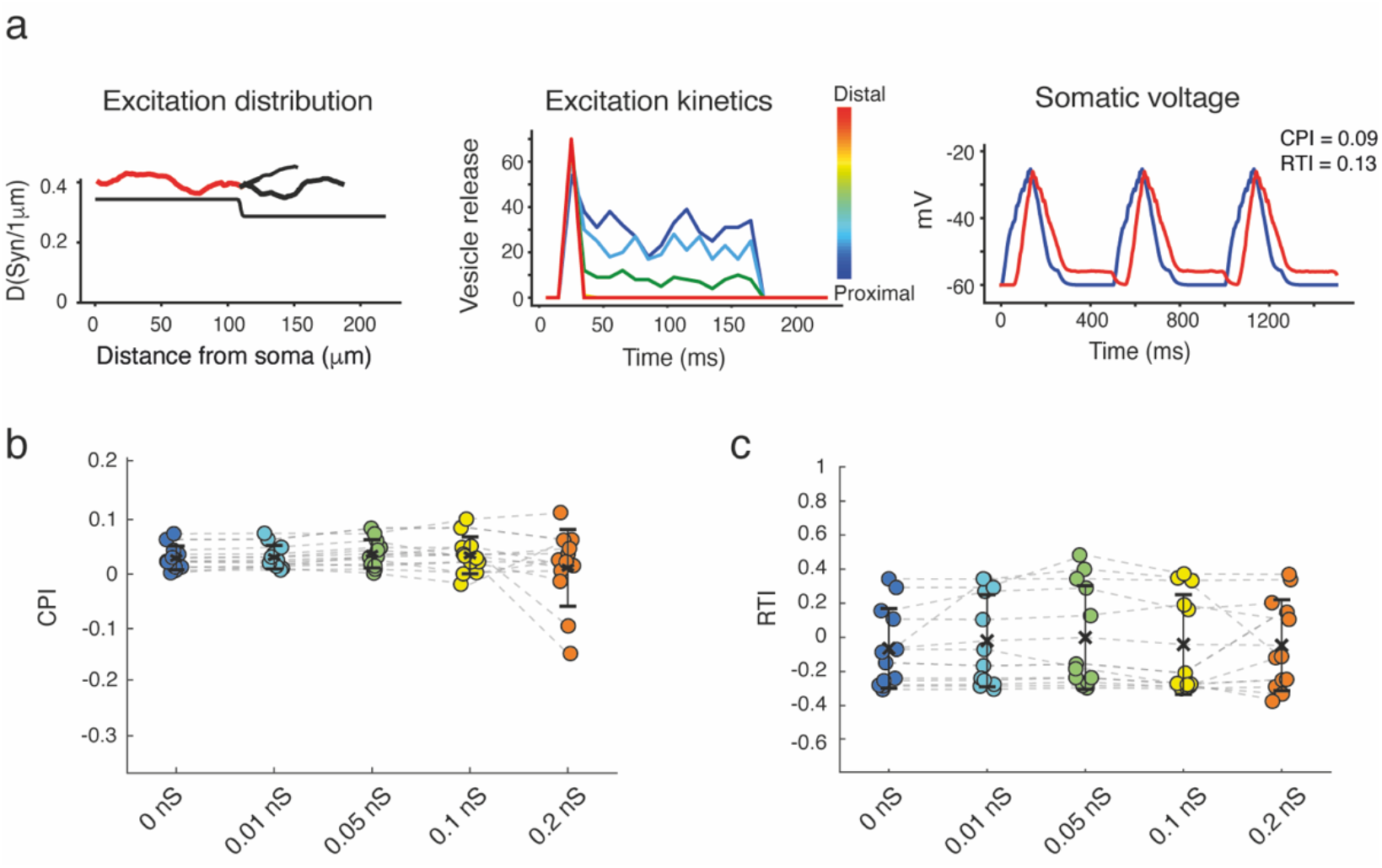
Inhibition is not sufficient to evoke centrifugal preference in non-CF preferring SACs. **a.** *Left*: Probability of excitatory synapses as a function of distance from soma for an example simulated SACs that did not display CF preference. *Center*: the color-coded kinetics of the simulated SAC excitatory inputs along the processes. *Right:* the voltage recorded from the soma of the simulated SAC in response to expanding (red) and collapsing (blue) rings. The CPI and RTI values are noted on the right. **b, c.** The CPI and RTI values of a population of non-CF preferring SACs (n=12).

**Supplementary Figure 9.**
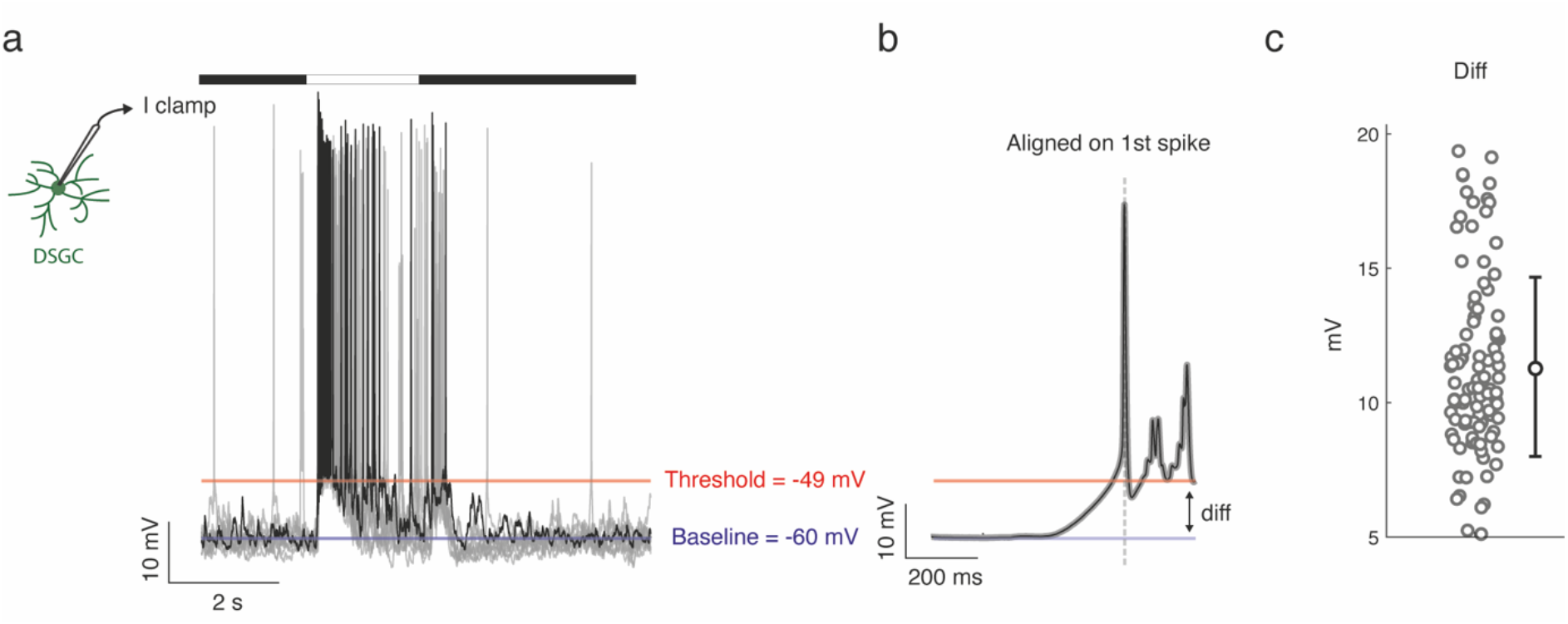
DSGC spiking properties. **a**. Example intracellular current-clamp recordings from a DSGC in response to presentation of a 2-sec bright spot (100 μm diameter). One repetition is shown in black, superimposed on nine other repetitions in grey. The Blue horizontal line denotes the baseline voltage of the cell (average of all traces), and the red line indicates the threshold for spiking. **b**. Average waveform of all ten repetitions, aligned to the peak of the first spike evoked in response to spots presentation. **c**. The difference between the threshold and the baseline voltage for all spikes elicited during a white spot presentation (11.3±3.3 mV; n = 154 spikes from 15 cells).

